# A highly potent, orally bioavailable pyrazole-derived cannabinoid CB2 receptor-selective full agonist for *in vivo* studies

**DOI:** 10.1101/2024.04.26.591311

**Authors:** Andrea Chicca, Daniel Batora, Christoph Ullmer, Antonello Caruso, Jürgen Fingerle, Thomas Hartung, Roland Degen, Matthias Müller, Uwe Grether, Pal Pacher, Jürg Gertsch

## Abstract

The cannabinoid CB2 receptor (CB2R) is a potential therapeutic target for distinct forms of tissue injury and inflammatory diseases. To thoroughly investigate the role of CB2R in pathophysiological conditions and for target validation *in vivo*, optimal pharmacological tool compounds are essential. Despite the sizable progress in the generation of potent and selective CB2R ligands, pharmacokinetic parameters are often neglected for *in vivo* studies. Here, we report the generation and characterization of a tetra-substituted pyrazole CB2R full agonist named RNB-61 with high potency (*K*_i_ 0.13–1.81 nM, depending on species) and a peripherally restricted action due to P-glycoprotein mediated efflux from the brain. ^3^H and ^14^C labelled RNB-61 showed apparent *K*_d_ values < 4 nM towards human CB2R in both cell and tissue experiments. The >6000-fold selectivity over CB1 receptors and negligible off-targets *in vitro*, combined with high oral bioavailability and suitable systemic pharmacokinetic (PK) properties, prompted the assessment of RNB-61 in a mouse ischemia-reperfusion model of acute kidney injury (AKI) and in a rat model of chronic kidney injury/inflammation and fibrosis (CKI) induced by unilateral ureteral obstruction. RNB-61 exerted dose-dependent nephroprotective and/or antifibrotic effects in the AKI/CKI models. Thus, RNB-61 is an optimal CB2R tool compound for preclinical *in vivo* studies with superior biophysical and PK properties over generally used CB2R ligands.

## Introduction

Cannabinoid CB1 receptors (CB1Rs) are expressed both in the central nervous system and peripherally and are responsible for the neuropharmacological effects of psychoactive cannabinoids like Δ9-THC^1,2^. In contrast, CB2 receptors (CB2Rs) are expressed primarily in the immune system and are responsible for few, if any, obvious behavioral effects^3–6^. The arachidonic acid-derived endocannabinoid lipids anandamide (AEA)^7^ and 2-arachidonoylglycerol (2-AG)^8^ non-selectively activate both CB receptors. Since endocannabinoids (eCBs) are rapidly degraded, metabolically stable agonists that selectively target CB1Rs and CB2Rs, respectively, have proven useful tools to elucidate their physiological roles and to modulate the endocannabinoid system (ECS)^9,10^. Importantly, the tissue protective role of CB2Rs in pathophysiological processes related to inflammation and their lack of central effects have rendered them an attractive drug target^11^. Consequently, structurally diverse CB2R-selective agonists are being developed as drug candidates^12^ for various diseases/pathological conditions ranging from chronic and inflammatory pain^13^, pruritus^14^, diabetic neuropathy^15^, liver cirrhosis^16^, and various types of ischemic-reperfusion injury^17–19^, to autoimmune, kidney and fibrotic diseases^4–6,15,20–26^. Although during the last two decades numerous selective and potent CB2R ligands belonging to diverse chemical scaffolds have been described in the scientific and patent literature, only a handful of synthetic ligands reached the clinical stage of development^12,27^. The reason for this may partly be attributed to the lack of knowledge regarding the different physiological roles of CB2Rs in cells and tissues. The use of conditional CB2R (*Cnr2*) knock-out mice significantly contributed to elucidate the role of CB2Rs in diverse pathophysiological conditions including liver and kidney inflammation and fibrosis^6,15,20,28,29^. Yet pharmacological probes bearing optimal pharmacokinetic (PK) properties represent a non-redundant complementary aid for target validation *in vivo*. An international consortium has previously profiled available CB2R ligands for basic research and concluded that JWH-133 and HU-308 were the best profiled CB2R agonists *in vivo*^30^. Nonetheless, due to their high lipophilicity, relatively low solubility, and strong binding to plasma proteins, these cannabinoid-like molecules show sub-optimal PK properties. Therefore, the generation and characterization of novel CB2R agonists combining high potency with a) high selectivity over other targets, especially the CB1R; b) favorable physicochemical properties including a balanced mixture of lipophilicity and water solubility; and c) good bioavailability, are crucial for pharmacological *in vivo* testing. Furthermore, the use of brain-impermeable ligands is desirable for peripheral indications such as acute or chronic kidney diseases.

In the absence of comparable and comprehensive *in vitro* assessments of promising CB2R scaffolds published in the scientific and patent literature, we synthesized five highly potent CB2R agonists: A-796260^31^ (**RNB-92**) and **RNB-61** from Abbott^32^, **RNB-73** from Amgen^33^ and **RNB-90**^34^ **and RNB-70**^35^ from the Boehringer Ingelheim. We characterized their CB2R binding affinity and selectivity over CB1R, as well as physiochemical and pharmacokinetic properties. **RNB-61**, which was not thoroughly profiled in the patent^32^, exhibited the highest selectivity towards CB2Rs and thus was profiled in-depth on its receptor pharmacology, bioavailability, PK and metabolism. Because numerous recent studies have demonstrated protective effects of CB2R signaling in various relevant preclinical models of acute and chronic kidney diseases (e.g. induced by ischemia/reperfusion (I/R), chemotherapy drug cisplatin, advanced liver injury (hepatorenal syndrome), chronic diabetes and unilateral ureteral obstruction (UUO)^15,20–26,29^, we also tested the efficacy of the compound in models of acute or chronic kidney injury/inflammation and/or fibrosis induced by renal I/R in mice or UUO in rats. Based on our data, we propose the pyrazole-derived **RNB-61** as an optimal pharmacological tool to interrogate the roles of peripheral CB2Rs *in vivo*.

## Materials and methods

### Synthesis of RNB compounds

The synthesis of **RNB-61**, **RNB-70**, **RNB-73**, **RNB-90**, and **RNB-92** were performed as described in the literature^31–35^. The synthesis of **RNB-61** and its regioisomer was accomplished as depicted in Scheme S1 and is described in more detail in the Supplementary Information. Briefly, the tosylate **(1)** was reacted with hydrazine hydrate and immediately converted in a [2+3] cycloaddition reaction to pyrazol **(2)**. Subsequent amide coupling using 2-fluoro-5-trifluoromethylbenzoyl chloride provided amide **(3)** in 66% yield. Regioselective methylation of the pyrazol N1 position using dimethyl sulfate generated tetra-substituted pyrazol **(4)** in 52% yield. Via nucleophilic aromatic substitution with 2-methylpropane-1,2-diol in the presence of potassium tert-butoxide the synthesis of **RNB-61** was finalized. Importantly, reaction of the amide with dimethyl sulfate in the presence of potassium carbonate led selectively to amide nitrogen methylation yielding the regioisomer of **RNB-61**.

### Synthesis procedures for radioligands

#### [^3^H]RNB-61

The *N*-desmethyl precursor of **RNB-61** (1.0 mg, 2.14 µmol) was added to a solution of [^3^H]methyl 4-nitrobenzenesulfonate (1.85 GBq, 0.160 mg, 0.714 µmol) in 50 µL of toluene (dried over aluminum oxide Woelm B Super I) in a screw-top vial and heated to 120 °C for 65 h. After evaporation of the solvent the crude product was purified by silica gel chromatography using a mixture of dichloromethane and methanol (95:5) as eluent. The isolated fractions were analyzed by radio-TLC on silica plates (dichloromethane/methanol/triethylamine, 90:10:1). The pure fractions were pooled, the solvent was removed under reduced pressure and the residue was dissolved in 10 mL of ethanol to yield 792 MBq (43%) of the tritium-labeled radioligand in a specific activity of 3.15 TBq/mmol (based on MS analysis) and 99.4% radiochemical purity (by radio-HPLC).

#### [^14^C]RNB-61

The *N*-desmethyl precursor of **RNB-61** (209 mg, 447 µmol) was added to a solution of [^14^C]methyl 4-nitrobenzenesulfonate (925 MBq, 97.8 mg, 446 µmol) in 1.5 mL of toluene (dried over aluminum oxide Woelm B Super I) in a screw-top vial and heated to 120 °C for 21 h. After evaporation of the solvent the crude product was purified by silica gel chromatography using a mixture of dichloromethane, methanol, and triethylamine (97:3:0.5) as eluent. The isolated fractions were analyzed by radio-TLC on silica plates (dichloromethane/methanol/triethylamine, 90:10:1). The pure fractions were pooled, and the solvent was removed under reduced pressure to yield 76.5 mg (307 MBq, 33%) of the ^14^C-labeled target compound as white solid in a specific activity of 1.94 GBq/mmol (based on MS analysis) and 99.2% radiochemical purity (by radio-HPLC).

### Receptor binding and activity assays

Competition and saturation binding assays were performed using the radiolabeled CB1R/CB2R agonist [^3^H]-CP55940 (PerkinElmer). Competition assays were conducted by incubating membrane protein fractions from human embryonic kidney (HEK) cells expressing the human CNR1 or CNR2 receptors with 1.5 nM [^3^H]-CP55940 in the presence or absence of increasing concentrations of **RNB-61** for 2 h at 30°C in a final volume of 0.2 mL of assay buffer (50 mmol L^-1^ Tris-HCl, 5 mmol L^-1^ MgCl_2_, 2.5 mmol L^-1^ EDTA, and 0.5% fatty acid-free BSA [pH 7.4] and 1% DMSO), with gentle shaking. Saturation binding assays were conducted by incubating membrane protein fractions from HEK cells with 12 concentrations in the range of 80–0.039 nM [^3^H]-CP55940 for 2 h at 30°C in a final volume of 0.2 mL/well of assay buffer without DMSO. WIN55212-2 (PerkinElmer) (10 µM) was used to define non-specific binding; >95% of the total binding signal was specific.

Binding reactions were terminated by vacuum filtration onto 0.5% polyethyleneimine presoaked GF/B filter plates (Packard) using a Filtermate cell harvester followed by 6 brief washes with 0.3 mL/well of ice-cold wash buffer. Wash buffer comprised 50 mmol L^-1^ Tris-HCl, 5 mmol L^-1^ MgCl_2_, 2.5 mmol L^-1^ EDTA, and 0.5% fatty acid-free BSA, pH 7.4. Plates were dried at 50°C for 1 h and liquid scintillation counting was used to determine levels of bound radiolabel. IC_50_ values and Hill slopes were determined with a 4-parameter logistic model using ActivityBase (ID Business Solution, Guilford, UK) and pKi values were determined using the Cheng-Prusoff equation. Binding data for 80 additional receptors was carried out at Eurofins Scientific (CEREP) and is reported as the average **RNB-61** induced percent inhibition of the binding of reference compounds in two measurements.

Functional CB2R activity was assessed with the cyclic AMP (cAMP) assay in Chinese hamster ovary (CHO) cells recombinantly expressing human wild type, human Q63R variant, cynomolgus, canine, rat and mouse CNR2, in CHO cells expressing human wild type CNR2 with the [^35^S]GTPγS assay and β-arrestin2 assay (Pathhunter assay, DiscoverX) as described previously^30^. Binding and functional assessment of endocannabinoid targets was done as depicted in our previous report^36^.

### Tissue radioligand experiments using [^3^H]RNB-61 and [^14^C]RNB-61

For the radioligand experiments, ten-week-old male C57BL/6J mice were obtained from the Jackson Laboratory (Bar Harbor, ME). Male Cnr2^−/−^ KO mice on C57BL/6J background and their wild-type controls (Cnr2^+/+^) were used in the study. The CB2R knockout allele was created by Deltagen. Specifically, the “Neo555T” construct was electroporated into 129P2/OlaHsd-derived E14 embryonic stem (ES) cells. Targeted mutant cell line 655 had homologous recombination of the “Neo555T” construct resulting in a 334 bp deletion in the coding exon of CB2R locus on chromosome 4. The donating investigator reported the *Cnr2^-^* mice were backcrossed at least five generations to C57BL/6J mice prior to sending to The Jackson Laboratory Repository. C57/black 6 mice or CB2R knock out mice were used in a model of LPS challenge. CB2R knock out mice on a C57/black 6 background were purchased from Jackson labs (B6.129P2-*Cnr2^tm1Dgen^*/J). The lipopolysaccharide (LPS) challenge was carried out by injecting 1 μg of LPS per mouse was i.p. 30 minutes after application of test compounds. 6h later mice were killed by cervical dislocation under deep anesthesia using Xylazine (10 mg kg^-1^) / Ketamine (100 mg kg^-1^). Spleens were removed and either used for membrane preparation or sliced on a cryostat at -20 °C. Slices were 20 µm thick, transferred to gelatin-coated slides and kept dry at -80 °C. For histological control, adjacent sections were stained with haematoxylin/eosin. Total crude spleen membranes were obtained following the method of Ploug et al. (1993). Frozen spleen samples were homogenized in in 210 mM sucrose, 40 mM NaCl, 2 mM ethylene glycol-bis (ß-aminoethylether) N,N,N’,N’-tetraacetic acid, 30 mM HEPES (pH 7.4), 0.35 mg ml^-1^ PMSF at pH 7.4; 10 mg ml^-1^) using a polytron homogenizer (Kinematica, Switzerland). Total spleen membranes were then recovered by centrifugation at 100 000 g at 40C for 90 min. The pellets were resuspended in 10μl/mg of 10 mM Tris–HCl and 1 mM EDTA (pH = 7.4), and then 4 μl mg^-1^ of 20% SDS was added. Samples were then centrifuged at 1100 g for 25 min. Protein concentrations of the supernatant was determined spectrophotometrically.

### Compound stability

The stability of the compound was assessed using the aqueous stability assay (ASTA), as previously described^37^. In short, aqueous solutions of **RNB-61** were prepared at 5 different pH values (range 1–10), added to incubation plates and shaken for 10 min at 37 °C. Solutions were transferred to a filter plate (Millipore MSGVN2250, pore size 0.22 μm) and filtered into V-bottom plates (ABGene, AB-0800) prior to heat-sealing. The procedure was repeated, increasing the 37 °C incubation time by 2 h. Samples were analyzed by HPLC at 0 and 2 h. A compound was classified as unstable if <90% of the initial concentration was detected after 2 h.

### Solubility and lipophilicity

For the determination of the octanol/water distribution coefficient (logD), the carrier-mediated distribution system (CAMDIS)-assay was used as described elsewhere^38^. Kinetic and thermodynamic solubility was assessed using the lyophilization solubility assay (LYSA) and thermodynamic solubility assay (THESA), respectively. For the LYSA, the solubility of RNB-61 in phosphate buffer at pH 6.5 from an evaporated 10 mM DMSO compound stock solution was measured. Two aliquots of the test compounds were dried and dissolved in phosphate buffer at pH 6.5. The solutions were then filtered and diluted (3 different dilution levels for each compound) before RapidFire MS analysis was performed. Each test compound was quantified using a 6-point calibration curve prepared with the same DMSO starting solution. For THESA, **RNB-61** (8.6 mg per mL solvent/vehicle) was stirred in HPLC vials (9 × 12 × 32 mm, Waters) at 350 rpm for 15 h. The presence of solid particles was determined by microscopic analysis of 10 μL samples. If the active pharmacological ingredient (API) was completely dissolved, more solid API was added and before stirring for another 15 h. This step was repeated for up to 96 h or until residual solid particles could be detected. Samples (0.5 mL) were transferred to Eppendorf Ultrafree filter tubes (Filter: PVDF 0.22 μm) and centrifuged at 14,500 rpm for 10 min. The filtrates were diluted in ethanol and analyzed by ultra-pure liquid chromatography (UPLC). Measurements were repeated in 0.05 M aqueous phosphate buffer, and in fasted (FaSSIF), and fed (FeSSIF) simulated gastrointestinal fluid.

### Hepatocyte and microsomal stability

The hepatocyte clearance assay was performed as previously described^39^. For animals, hepatocyte suspension cultures were either freshly prepared by liver perfusion studies or prepared from cryopreserved hepatocyte batches (pooled C57BL6 mouse hepatocytes were purchased from BioreclamationIVT (NY, USA)). For human, commercially available, pooled (5–20 donors), cryopreserved human hepatocytes from non-transplantable liver tissues were used. For the suspension cultures, Nunc U96 PP-0.5 mL (Nunc Natural, 267245) plates were used, which were incubated in a Thermo Forma incubator from Fischer Scientific (Wohlen, Switzerland) equipped with shakers from Variomag^®^ Teleshake shakers (Sterico, Wangen, Switzerland) for maintaining cell dispersion. The cell culture medium was William’s media supplemented with glutamine, antibiotics, insulin, dexamethasone and 10% FCS. Incubations of a test compound at 1 µM test concentration in suspension cultures of 1·10^6^ cells mL^-1^ (∼1 µg/µL protein concentration) were performed in 96-well plates and shaken at 900 rpm for up to 2 hours in a 5% CO_2_ atmosphere at 37 °C. After 3, 6, 10, 20, 40, 60, and 120 minutes, a 100 µL cell suspension in each well was quenched with 200 µL methanol containing an internal standard. Samples were then cooled and centrifuged before analysis by LC-MS/MS. Log peak area ratios (test compound peak area / internal standard peak area) or concentrations were plotted against incubation time with a linear fit. The slope of the fit was used to calculate the intrinsic clearance (CL_int_). Microsomal clearance data were generated as previously reported by our group^40^.

### Drug metabolism, CYP and hERG inhibition

Cytochrome P450 (CYP) assays were conducted as previously described^41^. GSH adduct formation data was assessed as reported in a previous study^42^. *h*ERG inhibition was measured as described in a recent publication^43^.

### Permeability assays

The general permeability of the compound was assessed using the parallel artificial membrane permeability (PAMPA) assay and the permeability-glycoprotein (P-gp) assay was used to specifically test for brain penetration. PAMPA data were generated as previously reported by our group^44^. P-gp (also known as multidrug resistance protein 1 (MDR1)) is the most studied and best characterized drug transporter. The P-gp assay evaluates the ability of test compounds to serve as a P-gp substrate. The assay was performed as described elsewhere^45^.

### Pharmacokinetics after intravenous and oral administration

To characterize the pharmacokinetic behavior of RNB-61, studies in rodents were conducted at the *in vivo* facility of F. Hoffmann-La Roche Ltd. (Basel, Switzerland). All animal experiments were performed in conformity with local animal welfare regulations for the care and use of laboratory animals.

The plasma-concentration time profile was studied in rodents after single-dose **RNB-61** administration by oral gavage (p.o. microsuspension) and intravenous bolus injection (i.v. solution). Three groups of rats (n=2 per group) were administered **RNB-61** either at 1 mg kg^-1^ i.v., 3 mg kg^-1^ p.o., or 26 mg kg^-1^ p.o.. Plasma samples were drawn at 0.083, 0.25, 0.5, 1, 2, 4, 8, and 24 post-dose in the i.v. group, and at 0.25, 0.5, 0.75, 1.5, 3, 5, 8, and 24 h post-dose in the p.o. groups. The samples were analysed for **RNB-61** concentration by LC/MS-MS. The pharmacokinetic parameters were estimated by non-compartmental analysis in the Certara WinNonlin software. A similar study design was run in mice with three groups receiving 2 mg kg^-1^ i.v., 4 mg kg^-1^ p.o., or 26 mg kg^-1^ p.o., and plasma sampling occurring in a composite profile (n=2 for each timepoint) up to 7 or 24 h post-dose.

Male C57/BL6 mice (n=5 per group) were used to study the influence of P-gp efflux on brain penetration. The mice were administered **RNB-61** by i.v. bolus injection at 1 mg kg^-1^ (2 groups). In order to block P-gp activity, an i.v. bolus of 5 mg kg^-1^ tariquidar^46^ was injected 30 minutes prior to **RNB-61**. Plasma was collected at 0.083, 0.25, 0.5, 2, 4, 7 h post-dose in a composite profile. Brain, CSF, and vitreous body were collected at terminal time points 0.5, 2, 4, 7 h post-dose.

### Kidney disease models

In the ischemia/reperfusion (I/R) model of acute kidney injury (AKI)^47,48^, the compounds were administered orally by gavage to C57BL/6 mice 30 minutes before ischemia obtained by clamping both renal arteries and veins for 25 minutes, followed by 24 hours of reperfusion. Mice were anesthetized using xylazine (10 mg kg^-1^) / ketamine (100 mg kg^-1^) injected intraperitoneally. Sham treated mice were treated identically except for the temporary closure of the renal vessels. After 24 hours, under re-anesthesia, plasma was taken for biomarker analysis. Thereafter, mice still in deep anesthesia were sacrificed by cervical dislocation. Creatinine, BUN (blood urea nitrogen), NGAL (neutrophil gelatinase-associated lipocalin), Osteopontin, and KIM1 (kidney injury molecule-1) levels were determined using commercially available standard assays. To assess fibrotic effects, we used the unilateral ureteral obstruction (UOO) model in Sprague Dawley rats^49^. For the UOO model, rats were sacrificed after 5, 8 and 11 days and the percentage of PicroSirius Red positive areas (indicator of collagen III-I deposition) were measured in 4 micrometer histological cross sections of the kidney after 2% paraformaldehyde perfusion fixation and paraffin embedding. The extend of kidney fibrosis was quantified based on PicroSirius Red staining of 10 renal sections about 1 mm apart from each kidney. The collagen-III-I-positive pixel counts were determined in the cortical aspect of each kidney section from 10 optical fields using a 40x objective. As the total area analyzed was identical for all kidneys under investigation, the absolute number of pixels was used as the relevant readout. Vehicle treated animals received saline gavage only. The details of PicroSirius Red staining were previously described^22^.

### Data analysis and statistics

All statistical analysis was carried out in Python 3.9. Data are represented as mean ± standard deviation unless otherwise indicated. Statistical significance was determined using Mann-Whitney test with the Bonferroni correction where applicable unless otherwise stated (^ns^p>0.05, *p<0.05, **p<0.01, ***p<0.001, ****p<0.0001).

## Results

### RBN-61 is a highly potent and selective CB2 receptor-specific ligand

With the aim to identify an optimal CB2R tool compound, we synthesized five highly potent CB2R agonist found in patent literature: A-796260^31^ (**RNB-92**) and **RNB-61** from Abbott^32^, **RNB-73** from Amgen^33^ and **RNB-90**^34^ and **RBN-70**^35^ from the Boehringer Ingelheim (Figure 1). The binding interactions of these CB2R agonists with cannabinoid receptors were assessed on both CB1Rs and CB2Rs, respectively (Table 1). Out of the five ligands, **RBN-61** potently bound to human CB2R (*h*CB2R) with an apparent *K*i value of 0.57 ± 0.03 nM and exhibited the best-in-class 6,800-fold selectivity over *h*CB1R (*K*i value = 4.3 μM) (Table 1). The regioisomer of **RNB-61** with the methylated amide (Figure 1) was inactive (data not shown). Similarly, the *K*i value of **RNB-61** for mouse CB2R (*m*CB2R) was 1.33 ± 0.47 nM, with a *m*CB2R/*h*CB2R K_i_ ratio of 2.3, which was slightly higher than for **RNB-73** (0.56). The range of K_i_ values for the selected ligands were comparable to the EC_50_ values obtained in the cAMP inhibition assays (Table 1). **RNB-61** also had favorable solubility, lipophilicity, clearance, and permeability in comparison to the four other molecules (Table 1.). In addition to CB1R, **RNB-61** also exhibited a great selectivity over the other endocannabinoid system (ECS) targets; lacking interactions up to 10 µM for FAAH, MAGL, ABHD6, ABHD12, endocannabinoid membrane transporter (EMT) and COX-2 (Table S1). The specificity of the compound against other ECS targets were evaluated functionally. **RNB-61** did not inhibit the hydrolysis (FAAH, MAGL and ABHDs), oxygenation (COX-2) and cellular AEA uptake (EMT) of ECs at the screening concentration of 10 μM (Table S1). Next, a thorough characterization of **RBN-61** receptor pharmacology against 80 additional receptors, including dopamine, serotonin, epinephrine, norepinephrine, acetylcholine, GABA, benzodiazepine, opioid, adenosine, prostaglandin, and chemokine receptors, among others, was conducted using CEREP (now Eurofins). **RNB-61** showed no significant binding (defined as ≥ 50% of probe displacement) to most receptors at the screening concentration of 10 μM (Figure S1). The one exception was the Na^+^ ion channel site 2, for which 10 μM of **RNB-61** inhibited 96% of [^3^H]-batrachotoxin binding. Follow-up measurements with 1 μM and 0.1 μM of **RNB-61** showed 32% and no inhibition of [^3^H]-batrachotoxin binding, respectively, indicating no potential off-target interactions at physiologically relevant concentrations (<1 µM), thus supporting the very high selectivity of **RBN-61** towards CB2R (Figure S1).

**Figure 1:**
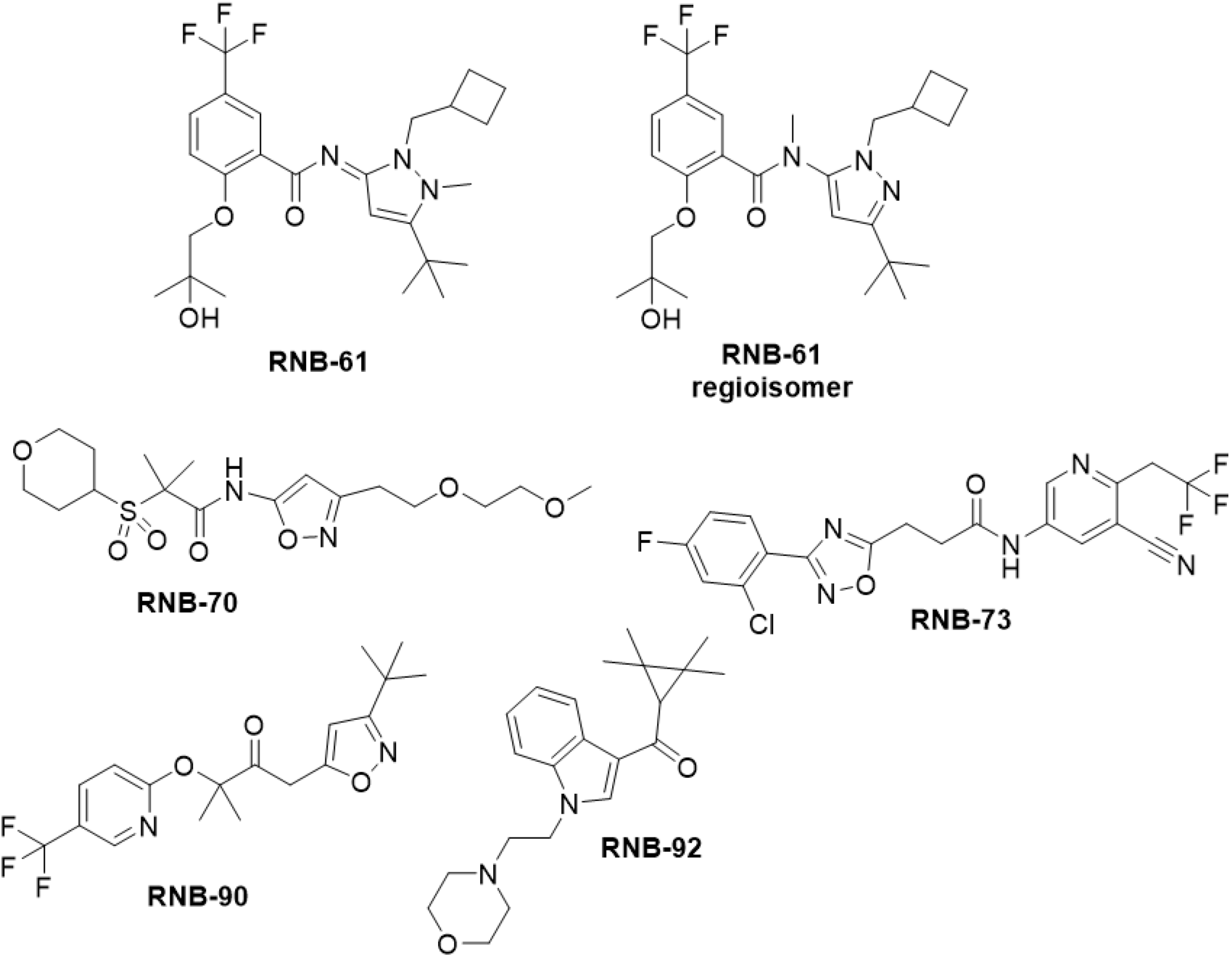
Chemical Structures of RNB compounds.

**Table 1:**
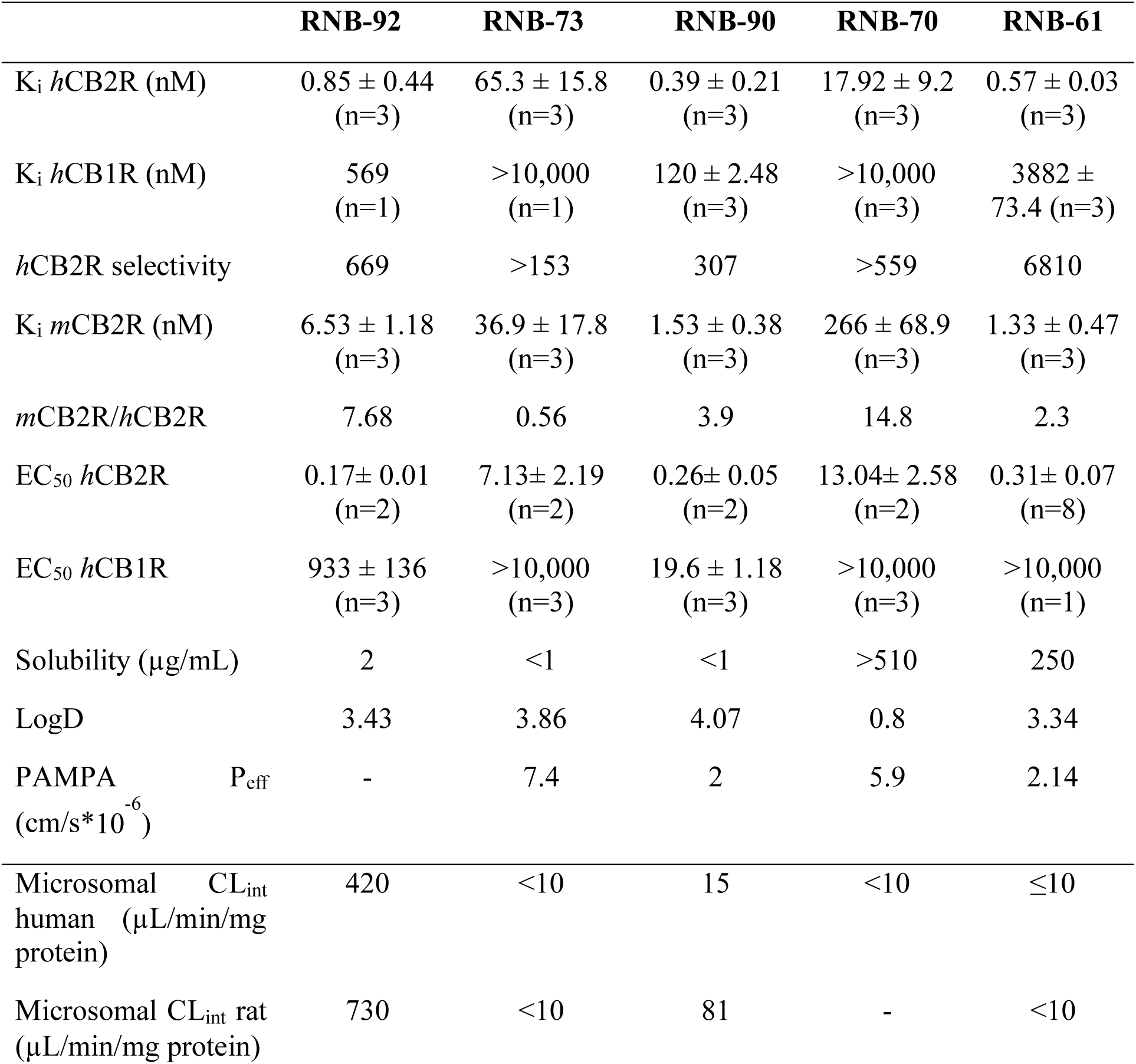
Comparison of the selectivity towards human CB2 (*h*CB2) receptors of five representative molecular scaffolds.

|  | **RNB-92** | **RNB-73** | **RNB-90** | **RNB-70** | **RNB-61** |
| --- | --- | --- | --- | --- | --- |
| K<sub>i</sub> *hCB2R* (nM) | 0.85 ± 0.44<br>(n=3) | 65.3 ± 15.8<br>(n=3) | 0.39 ± 0.21<br>(n=3) | 17.92 ± 9.2<br>(n=3) | 0.57 ± 0.03<br>(n=3) |
| K<sub>i</sub> *hCB1R* (nM) | 569<br>(n=1) | >10,000<br>(n=1) | 120 ± 2.48<br>(n=3) | >10,000<br>(n=3) | 3882 ±<br>73.4 (n=3) |
| *hCB2R* selectivity | 669 | >153 | 307 | >559 | 6810 |
| K<sub>i</sub> *mCB2R* (nM) | 6.53 ± 1.18<br>(n=3) | 36.9 ± 17.8<br>(n=3) | 1.53 ± 0.38<br>(n=3) | 266 ± 68.9<br>(n=3) | 1.33 ± 0.47<br>(n=3) |
| *mCB2R/hCB2R* | 7.68 | 0.56 | 3.9 | 14.8 | 2.3 |
| EC<sub>50</sub> *hCB2R* | 0.17± 0.01<br>(n=2) | 7.13± 2.19<br>(n=2) | 0.26± 0.05<br>(n=2) | 13.04± 2.58<br>(n=2) | 0.31± 0.07<br>(n=8) |
| EC<sub>50</sub> *hCB1R* | 933 ± 136<br>(n=3) | >10,000<br>(n=3) | 19.6 ± 1.18<br>(n=3) | >10,000<br>(n=3) | >10,000<br>(n=1) |
| Solubility (µg/mL) | 2 | <1 | <1 | >510 | 250 |
| LogD | 3.43 | 3.86 | 4.07 | 0.8 | 3.34 |
| PAMPA P<sub>eff</sub><br>(cm/s*10<sup>-6</sup>) | - | 7.4 | 2 | 5.9 | 2.14 |
| Microsomal CL<sub>int</sub><br>human (µL/min/mg<br>protein) | 420 | <10 | 15 | <10 | ≤10 |
| Microsomal CL<sub>int</sub> rat<br>(µL/min/mg protein) | 730 | <10 | 81 | - | <10 |

### RBN-61 is a CB2 receptor full agonist

A further characterization of the functional activity of **RBN-61** on *h*CB2R was performed measuring G-protein activation ([^35^S]GTPγS assay), cAMP formation and β-arrestin2 recruitment. As expected, **RBN-61** induced a concentration-dependent increase of [^35^S]GTPγS binding (EC_50_ = 0.33±0.09 nM) and β-arrestin2 recruitment (EC_50_ = 13.3±1.9 nM), while it inhibited forskolin (FSK)-driven cAMP formation (EC_50_ = 1.65±0.96 nM) (Figure 2). The functional modulation of CB2R activity occurred at a similar concentration range as the K_i_ and K_d_ value calculated for the binding. **RBN-61** induced the same maximal effect as CP55,940 for the inhibition of cAMP formation and [^35^S]GTPγS binding, while reaching approximately 80% of total β-arrestin2 recruitment (Figure 2). To decipher whether RNB-61 is suitable for the elucidation of the pharmacology of CB2R in different animal disease models, we next performed the cAMP assay with CB2R from species of preclinical interest as well as the common human missense variant CAA/CGG (Q63R). As shown in Table 2, **RNB-61** inhibited the FSK-induced cAMP formation on mouse, rat, dog, and monkey CB2Rs with EC_50_ values ranging from 0.13 to 1.86 nM, which were in line with the potency towards the wild type *h*B2 receptor (0.31±0.07 nM). Similarly, the EC_50_ value for the Q63R human variant was 0.29 ±0.05 nM.

**Figure 2:**
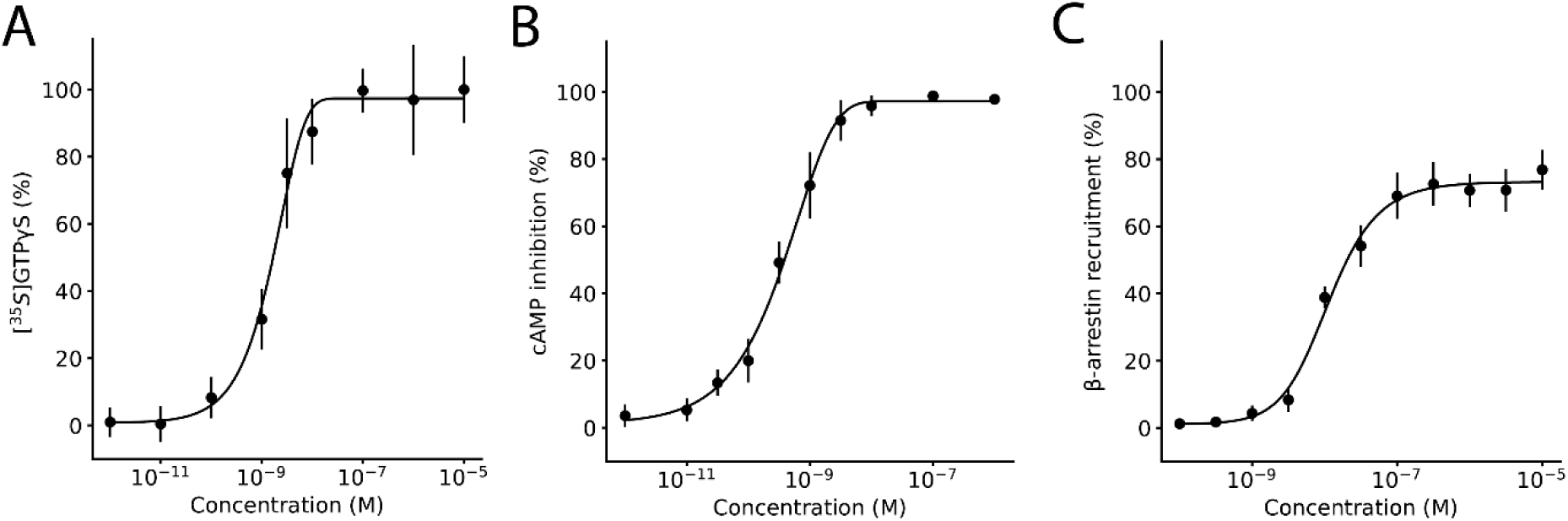
Concentration-response of RNB-61 on [^35^S]GTPγS binding cAMP inhibition and β-arrestin recruitment. A: Concentration-response of **RNB-61** on G-protein coupled receptor activation expressed as the percentage of binding of the non-hydrolysable GRP analog [^35^S]GTPγS (n=4, in triplicates, EC_50_=1.65±0.96 nM, E_max_ = 100) B: Concentration-response of **RNB-61** on cyclic AMP (cAMP) inhibition (n=6, in triplicates, EC_50_=0.33±0.09 nM, E_max_=100) C: Concentration-response of **RNB-61** on β-arrestin2 recruitment expressed as a percentage, normalized to 1 µM of CP55940 (n=6, in triplicates, EC_50_=13.3±1.9 nM, E_max_=80)

**Table 2:**
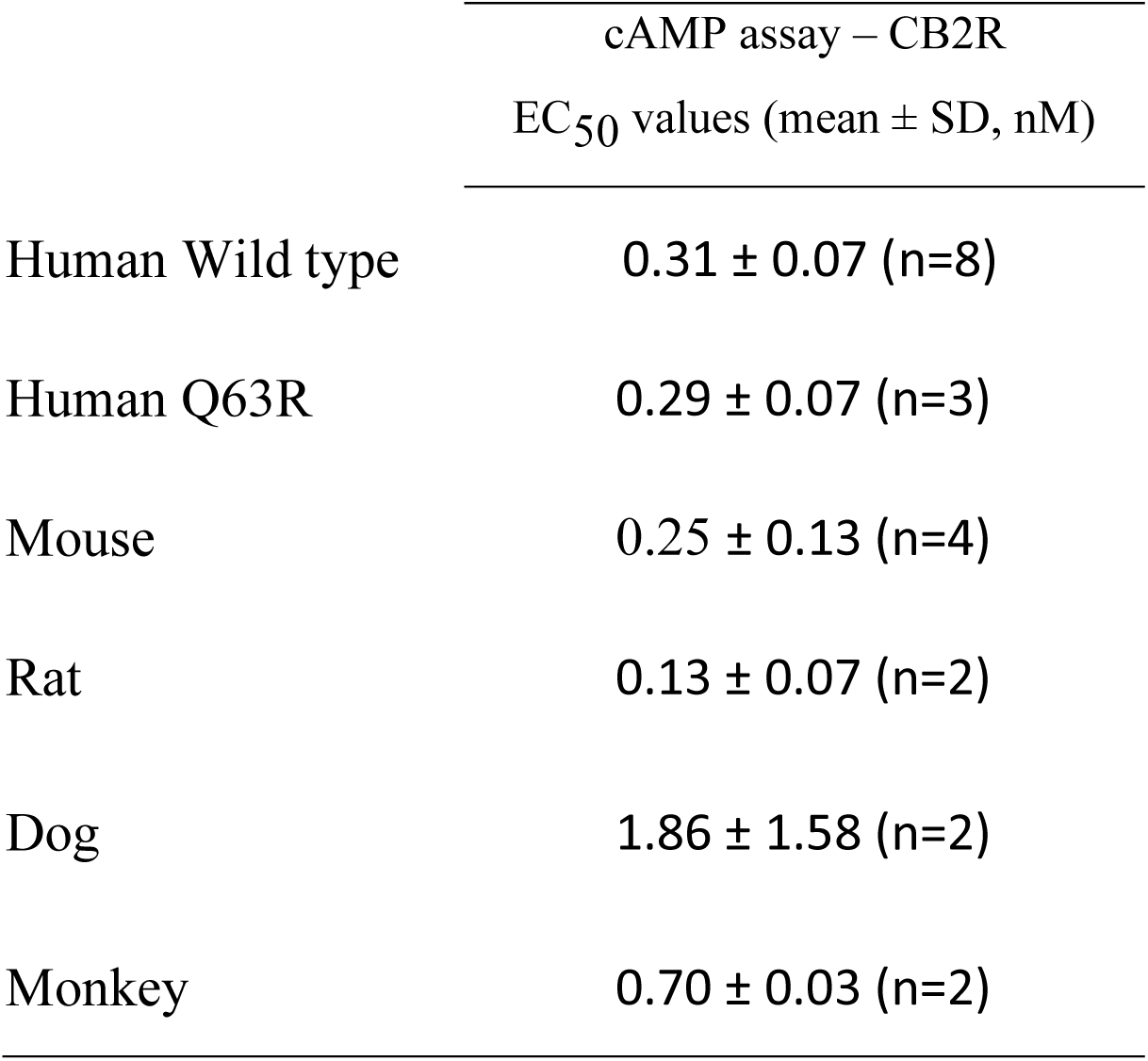
Comparison of EC_50_ values on cAMP production expressing CB2 from different species and the human Q63R mutant.

### [^3^H]-RNB-61 and [^14^C]-RNB-61 as radiopharmaceutical tools

Next, we validated radiolabeled versions of **RNB-61** by labeling CB2R expressing membrane preparations from cells and intact tissues. In labeling membrane preparation of CHO cells overexpressing the *h*CB2 receptors, [^3^H]-RNB-61 exhibited a K_d_ value of 3.08±0.61 nM, whereas [^14^C]-RNB-61 showed a K_d_ value of 3.62±2.31 nM, both in range of the binding interaction data (Figure 3). No non-specific labeling was observed below 50 nM for [^3^H]-**RNB-61** with a corresponding signal-to-background value of 6.0 at 250 nM, whereas for [^14^C]-**RNB-61**, the signal-to-background value was 1.6 at 250 nM. Next, we labeled *ex vivo* spleen slices from mouse with 200 nM of [^14^C]-**RNB-61**, which showed a prominent increase in radioactivity that could be competed with 10 µM of the non-selective CBR agonist, WIN 55,212-2 and was not detected in a CB2R KO mouse strain (Figure 3). We further elucidated the CB2R specificity of [^3^H]-**RNB-61** on spleen membranes in both rats and mice. In wild type rat and mice spleen membranes, the radioactivity could be competed away with both 10 µM WIN 55,212-2 and 1 µM of the CB2R specific antagonist SR144528 (SR2), however no significant reduction in the radioactivity was observed with 1 µM the CB1R specific antagonist SR141716A (SR1 or rimonabant) (Figure 3). Next, we evaluated the utility of **RNB-61** for CB2R expression in 17 different cell lines that included Chinese hamster ovary (CHO) cells that recombinantly express *h*CB1R or *h*CB2R, immune cells (HL-60, U937, Jurkat, RAW264.7, HMC-1, BV2), neuronal cells (SH-SY5Y, Neuro2a, NT18G2), and miscellaneous cell lines (HeLa, PC-12, HEK-293, HPKV). Out of the 17 cell lines, the *h*CB2R-overexpressing CHO cells, and all immune cells except for HMC-1 (P value=0.31) showed a statistically significant difference in radioactivity between the vehicle and the WIN55,212-2 preincubated samples (Figure 3). To confirm the differential CB2R radiolabeling induced in inflammatory conditions, we measured CB2R expression using [^14^C]-**RNB-61** in *ex vivo* spleen slices from mice challenged with lipopolysaccharide (LPS). We detected a significant increase in radioactivity after 6-24 hours of the introduction of LPS in the spleen which was completely absent in the CB2R KO mouse strain (Figure 3).

**Figure 3:**
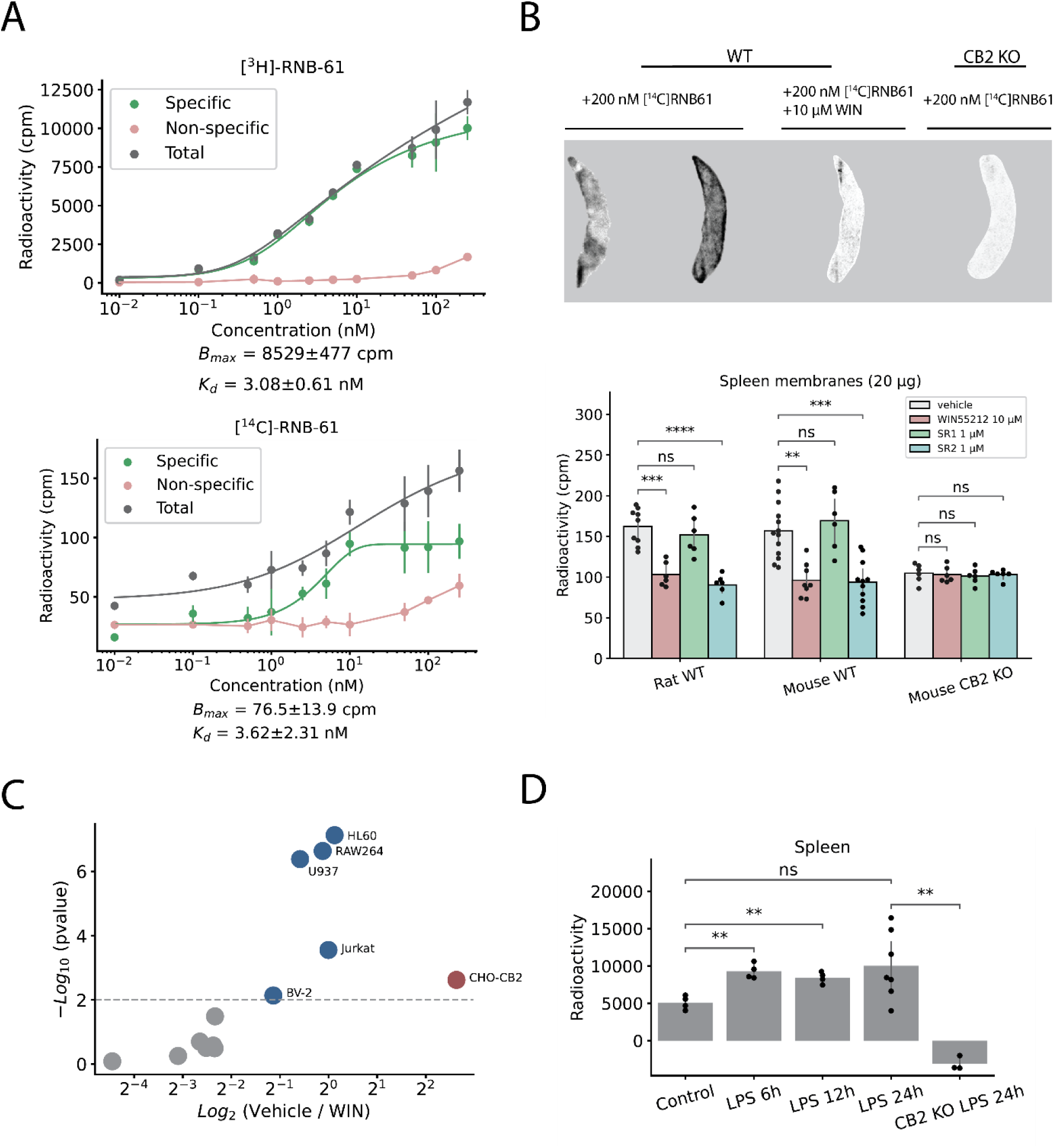
[^3^H]-RNB-61 and [^14^C]-RNB-61 to label CB2 receptors. A: Concentration-response of membrane binding of **[^3^H]-RNB-61** and **[^14^C]-RNB-61** on CHO cells recombinantly expressing the human CB2R (*h*CB2R) (n=4-6, B_max_=8529±477 cpm, K_d_=3.08±0.61 nM for **[^3^H]-RNB-61** and B_max_=76.5±13.9 cpm, K_d_=3.62±2.31 nM for **[^14^C]-RNB-61**). The non-specific binding was calculated by pre-incubating the cells with 10 µM of WIN 55,212-2. B: The top image depicts 4 representative images of spleen slices incubated in 200 nM of **[^14^C] RNB-61**. Images were acquired using the Typhoon FLA 9500 imaging system. The bottom plot depicts radioactivity of spleen membrane preparations from either wild-type (WT) rat, WT mouse or the CB2R knock-out mouse strain (n=6-13). In the WT rat and mouse, addition of 10 µM of the non-specific CBR agonist WIN55,212-2 resulted in a significant reduction in the detected radioactivity in cpm (p<0.01, p<0.001 for WT rat and WT mouse, respectively, independent t-test). Addition of 1 µM of the CB1R selective SR1 did not result in a significant difference, whereas the addition of 1 µM of the CB2R selective SR2 significantly reduced radioactivity (p<0.001, p<0.0001 for WT rat and WT mouse, respectively, independent t-test). No difference in radioactivity was observed for the CB2R KO mice in any condition. C: 17 different cell lines (CHO cells, immune cells, neuronal cells, miscellaneous cells) were incubated with 1 nM of **[^3^H]-RNB-61** and either vehicle or 1 µM of WIN55,212-2. The volcano-plot depicts the difference between the vehicle and the WIN-treated cells on the x axis and the negative logarithm of the p-value on the y axis. Only the *h*CB2 overexpressing CHO cells and immune cells were statistically significant in CB2R specific labeling (p<0.01). No significant labeling was detected in any of the neuronal cell lines (SHSY5Y, Neuro2a, NT18G2). D: CB2R expression measured in cpm in *ex vivo* spleen slices from mice challenged with LPS detected with the administration of 200 nM **[^14^C]RNB-61**. Bar chart depicts the specific signal which was attained by subtracting the non-specific signal (10 µM of WIN-55,212 co-incubation). LPS induced a statistically significant increase in CB2R expression at various timepoints (6-24h), which was completely absent in the CB2R KO mice.

### Physicochemical profile and stability

The physicochemical profile and stability of **RNB-61** were further evaluated in detail (Table 3). **RNB-61** showed a low partition coefficient (logD = 3.3), indicating moderate lipophilicity, which accounts for a good balance between solubility and permeability. Accordingly, **RNB-61** was stable in aqueous solution for 2 h at 37 °C at different pH values (1, 4, 6.5, 8 and 10) and showed a high solubility in four different assays (LYSA = 194 μg ml^-1^; THESA = 316 μg ml^-1^; FaSSIF = 630 μg ml^-1^; FeSSIF = 1373 μg ml^-1^). In the PAMPA assay, RNB-61 permeated through the cell membrane with an effective permeability (P_eff_) of 2.14 x 10^-6^ cm s^-1^, indicating a relevant permeability in cellular membranes. An estimation of the blood-brain-barrier penetration was performed *in vitro* using human and mouse P-glycoprotein overexpressing systems. The results showed a remarkably high P-gp efflux ratio (ER) suggesting that **RNB-61** is a strong P-gp substrate and might not reach bioactive concentration in the CNS. In further *in vitro* studies the metabolic stability of **RNB-61** was assessed, showing a low intrinsic clearance in purified microsomes and hepatocytes. Furthermore, the ligand did not affect the activity of the three most relevant cytochrome P450 isoenzymes 2C9, 2D6, 3A4 (IC_50_ value > 50 μM). Similarly, **RNB-61** did not form any adducts with glutathione in human nor in rat liver microsomes and it did not interact with the *h*ERG channel up to a concentration of 10 μM (Table 3). Overall, **RNB-61** showed a favorable balance between lipophilicity and solubility associated with optimal stability *in vitro*.

**Table 3:** Physicochemical and ADMET properties of RNB-61.

|  |  |
| --- | --- |
| Molecular weight | 481.556 |
| pKa | 7.33 |
| clogP | 3.34 (logD) |
| Polar surface area | 51 Å<sup>2</sup> |
| Hydrogen bond donors | 1 |
| Stability in aqueous solution<br>at pH 1, 4, 6.5, 8, 10 | 104.4, 102.7, 102.7, 100.5, 101.6<br>% of initial [200 nM] |
| Aqueous solution<br>LYSA, THESA, FaSSIF, FeSSIF | 194, 316, 630, 1373 µg mL<sup>-1</sup> |
| Melting poing | 188 °C |
| PAMPA P<sub>eff</sub> (%Acc/%Mem/%Don) | 2.14 x 10<sup>-6</sup> cm s<sup>-1</sup> (6/72/22) |
| P-gp transporter ER human | 40.6 |
| P-gp transporter ER mouse | 26.8 |
| Microsomal CL<sub>int</sub> human, rat, mouse | ≤10 µL min<sup>-1</sup> (mg protein)<sup>-1</sup> |
| Hepatocyte CL<sub>int</sub> human, mouse | 85, 48.5 µL min<sup>-1</sup> (10<sup>6</sup> cells)<sup>-1</sup> |
| IC<sub>50</sub> CYP450 3A4, 2C9, 2D6 inhibition | 35.5, 50, 15.5 µM |
| GSH (human liver microsomes) | no adducts detected |
| hERG inhibition | >10 µM |
| Fraction unbound in plasma (human, rat, mouse) | 1.9, 1.7, 1.9% |

### RNB-61 pharmacokinetics and brain penetration

The absorption and disposition of **RNB-61** was assessed in single-dose PK studies in rodents. Upon i.v. injection of 1 mg kg^-1^ bolus in rats, **RNB-61** exhibited a low plasma clearance (CL= 3.5 mL min^-1^ kg^-1^), in agreement with the *in vitro* CL_int_ values, and an intermediate volume of distribution (V_ss_=1.6 L kg^-1^), resulting in a terminal half-life of 6.0 h (Figure 4, Table 4). After oral administration of **RNB-61** 3 and 26 mg kg^-1^, the compound displayed high bioavailability, suggesting nearly complete absorption in the tested dose range. The maximal plasma exposure for 3 and 26 mg kg^-1^ p.o. dosing was reached at 5.5 and 3.0 h post-injection, respectively, with C_max_ values of 446 and 1710 ng mL^-1^. Taken together, the pharmacokinetic results indicate that in rats, RNB-61 reaches high nanomolar to micromolar concentrations in plasma after single-dose administration (C_max_= 1285 nM, 926 nM, 3.5 μM after 1 mg kg^-1^ i.v., 3 mg kg^-1^ p.o., 26 mg kg^-1^ p.o., respectively). The overall pharmacokinetic profile was similar in mice, as depicted in Figure 4 and Table 4.

**Figure 4:**
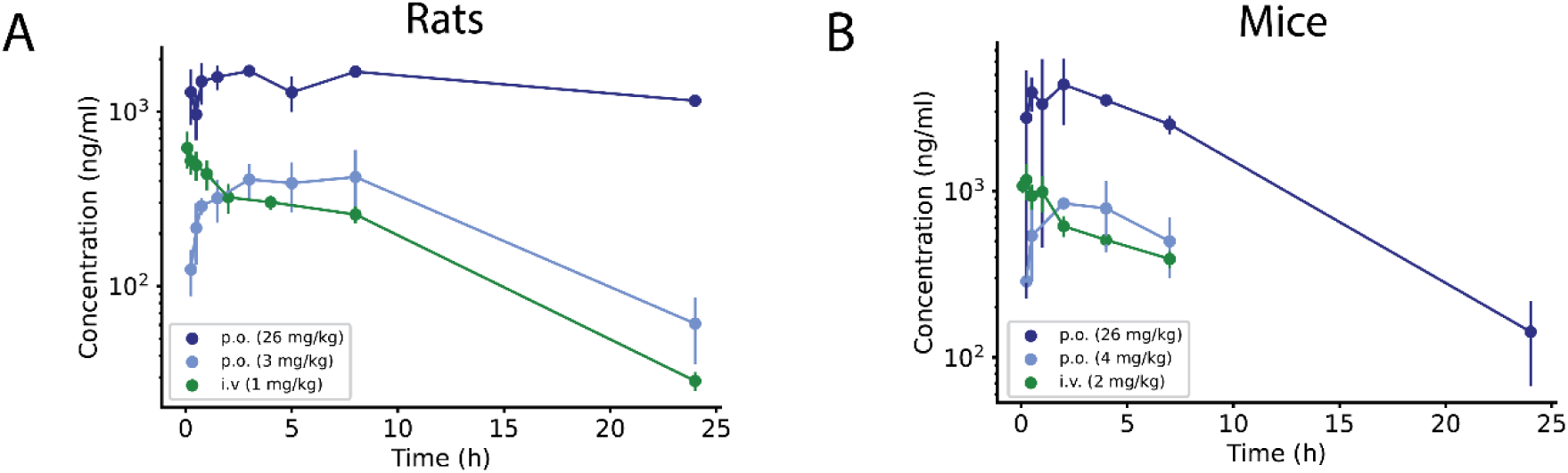
RNB-61 plasma pharmacokinetics after intravenous and oral administration. Plasma concentration (mean and standard deviation) of **RNB-61** after p.o. and i.v. bolus administration in (A) rats and (B) mice (n=2 for each timepoint).

**Table 4:** Pharmacokinetic parameters of RNB-61 in rodent models.

| **Rats** | **1 mg kg<sup>-1</sup>** | **3 mg kg<sup>-1</sup>** | **26 mg kg<sup>-1</sup>** |
| --- | --- | --- | --- |
| Route of Administration | i.v. | p.o. | p.o. |
| C<sub>max</sub> (ng mL<sup>-1</sup>) | 619 | 446 | 1710 |
| T<sub>max</sub> (h) | 0.083 | 5.5 | 3 |
| AUC<sub>inf</sub> (h ng mL<sup>-1</sup>) | 5173 | 7406 | 117100 |
| CL (mL min<sup>-1</sup> kg<sup>-1</sup>) | 3.5 | n/a | n/a |
| V<sub>ss</sub> (L kg<sup>-1</sup>) | 1.6 | n/a | n/a |
| T<sub>1/2</sub> (h) | 6.0 | 7.1 | 49 |
| Bioavailability (%) | n/a | 48 | 87 |

**Table 4: Pharmacokinetic parameters of RNB-61 in rodent models**
| **Mice** | **2 mg kg<sup>-1</sup>** | **4 mg kg<sup>-1</sup>** | **26 mg kg<sup>-1</sup>** |
| --- | --- | --- | --- |
| Route of Administration | i.v. | p.o. | p.o. |
| C<sub>max</sub> (ng mL<sup>-1</sup>) | 1167 | 843 | 4370 |
| T<sub>max</sub> (h) | 0.25 | 2 | 2 |
| AUC<sub>inf</sub> (h ng mL<sup>-1</sup>) | 8632 | 9300 | 47320 |
| CL (mL min<sup>-1</sup> kg<sup>-1</sup>) | 3.9 | n/a | n/a |
| V<sub>ss</sub> (L kg<sup>-1</sup>) | 2.4 | n/a | n/a |
| T<sub>1/2</sub> (h) | 7.7 | 6.4 | 4.3 |
| Bioavailability (%) | n/a | 54 | 42 |

Next, we investigated the brain exposure to **RNB-61**. Upon i.v. injection of 1 mg kg^-1^, **RNB-61** reached the peak brain concentration of 41.3 ng mL^-1^, which was >20-fold lower compared to the plasma level (C_max_= 972 ng mL^-1^) (Table 5). Similarly, the area under the curve from time zero to the last measurable concentration (AUC_last_) was >10-fold lower in the brain compared to plasma (AUC_last_= 215 vs. 2429 h * ng mL^-1^). The brain-plasma partition coefficient (Kp, brain) of **RNB-61**, calculated as the ratio of AUC_last_ in brain and plasma, was accordingly very low (Kp= 0.088), indicating a negligible penetration into the brain. In agreement with data obtained *in vitro* showing that **RNB-61** is a P-gp substrate, upon injection of the P-gp inhibitor tariquidar (5 mg kg^-1^), **RNB-61** reached a higher brain exposure (C_max_= 342 ng mL^-1^, AUC_last_ = 1877 h * ng mL^-1^). Both parameters were ∼10-fold higher than the corresponding values after injection of **RNB-61** alone, with no apparent differences in the plasma PK profile. The significantly increased penetration into the brain was confirmed by the Kp value of 0.747 (Table 5).

**Table 5:** Brain exposure of RNB-61 in male C57/BL6 mice (n=5/group) is enhanced by co-administration of tariquidar, a P-gp inhibitor. *Abbreviations: Kp,* brain-plasma partition coefficient, calculated as the ratio of AUC_last_ between brain and plasma.

|  |  | Plasma |  |  |  |  |  |
| --- | --- | --- | --- | --- | --- | --- | --- |
|  |  | T<sub>1/2</sub> | T<sub>max</sub> | C<sub>max</sub> | AUC<sub>last</sub> | AUC<sub>inf</sub> | CL |
| RNB-61 | Tariquidar | h | h | ng/mL | h*ng/mL | h*ng/mL | mL/h/kg |
| 1 mg kg<sup>-1</sup> | - | 4.00 | 0.25 | 972 | 2429 | 3318 | 301 |
| 1 mg kg<sup>-1</sup> | 5 mg kg<sup>-1</sup> | 3.68 | 0.083 | 737 | 2514 | 3558 | 281 |

**Table 5: Brain exposure of RNB-61 in male C57/BL6 mice (n=5/group) is enhanced by co-administration of tariquidar, a P-gp inhibitor.**
|  |  | Brain |  |  |  |  |  |
| --- | --- | --- | --- | --- | --- | --- | --- |
|  |  | T<sub>1/2</sub> | T<sub>max</sub> | C<sub>max</sub> | AUC<sub>last</sub> | AUC<sub>inf</sub> | K<sub>p</sub> |
| RNB-61 | Tariquidar | h | h | ng/mL | h*ng/mL | h*ng/mL | - |
| 1 mg kg<sup>-1</sup> | - | 5.98 | 0.58 | 41.3 | 215 | 375 | 0.088 |
| 1 mg kg<sup>-1</sup> | 5 mg kg<sup>-1</sup> | 8.42 | 0.53 | 342 | 1877 | 4386 | 0.747 |

### RNB-61 exerts protective effects in kidney ischemia reperfusion (I/R) and unilateral ureteral obstruction (UUO) rodent models

Based on its high potency and selectivity as CB2R agonist, together with the favorable PK profile, **RNB-61** represented a suitable tool compound to further investigate CB2R pharmacology *in vivo*. To validate the efficacy of **RNB-61**, evaluated its effects in two rodent models of kidney injury: the I/R-induced acute kidney injury (AKI) (Figure 5) and the UUO-induced model of chronic kidney injury (CKI), inflammation and progressive renal fibrosis (Figure 6). In the I/R model, we observed an over 2-fold, significant increase (1.19±0.33 mg kg^-1^) in plasma creatinine levels compared to the sham controls (0.55±0.09 mg kg^-1^), consistent with AKI (p<0.001, independent t-test). Similarly, blood urea nitrogen (BUN) levels increased from 26.6±5.6 mg dl^-1^ in the sham control to 49.8±8.9 mg dl^-1^ in the vehicle (p<0.001, independent t-test).The I/R-induced increases in creatinine and BUN plasma levels were hampered by the pre-treatment with **RNB-61** in a dose-dependent manner (Figure 5). The maximal protection was achieved at the dose of 3 mg kg^-1^ (48% and 30% reduction of creatinine and BUN compared to vehicle, respectively) similarly to the positive control fenoldopam at 20 mg kg^-1^ (creatinine and BUN levels reduced by 53% and 39% compared to vehicle, respectively) (Figure 5). In the same model, the AKI biomarkers neutrophil gelatinase-associated lipocalin (NGAL), kidney injury molecule-1 (KIM-1) and osteopontin were measured in plasma. In line with the effects observed on creatinine and BUN levels, **RNB-61** dose-dependently inhibited the release of all three biomarkers starting from 0.3 mg kg^-1^ (NGAL and KIM-1) and 3 mg kg^-1^ (osteopontin) and reaching the maximal protection at 3–30 mg kg^-1^ (reduction by 39%, 43% and 50% compared to vehicle for NGAL, osteopontin and KIM-1, respectively) similarly to the positive control tempol (at the dose of 50 mg kg^-1^, reduction by 43%, 49% and 57% compared to vehicle for NGAL, osteopontin and KIM-1, respectively). The nephroprotective effect of **RNB-61** was further evaluated in the UUO-induced model of kidney fibrosis in rats. Initial experiments were performed using the positive control, enalapril (32 mg kg^-1^ day^-1^) or vehicle to identify the best time to assess the anti-fibrotic effect after ureteral obstruction. A significant anti-fibrotic effect was evident on day 8 (55% reduction vs. vehicle) without any further improvement on day 11 (57% reduction vs. vehicle). Thus, 8 days were chosen as a suitable duration to assess the anti-fibrotic effect of **RNB-61**. As shown in Figure 6, **RNB-61** exerted potent antifibrotic effects in the full range of tested doses (0.3–10 mg mL^-1^). At 3 mg kg^-1^, it inhibited the accumulation of collagen-III-I by 61%, which was comparable to the protective effect of the positive control enalapril (inhibition by 52%).

**Figure 5:**
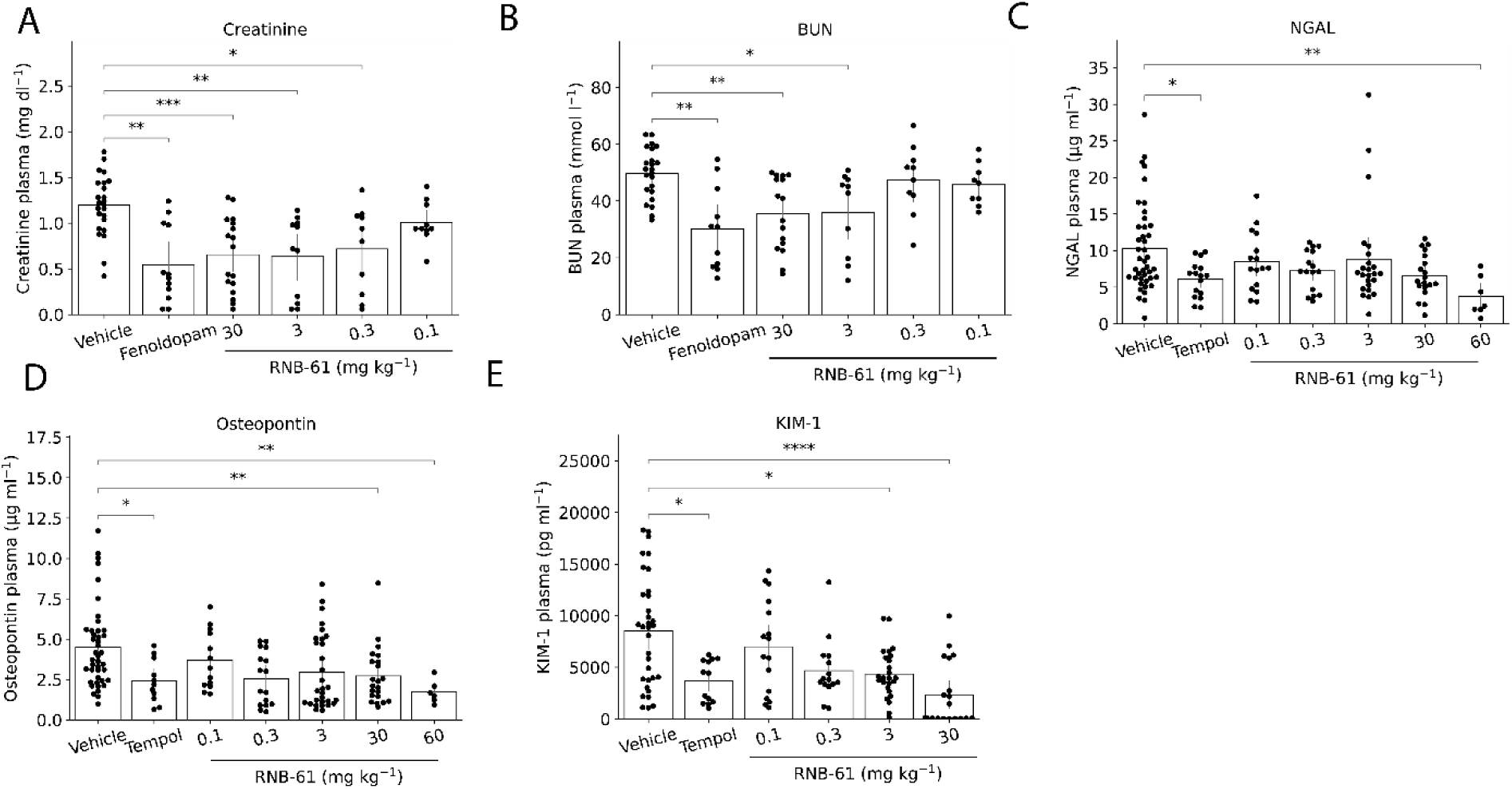
RNB-61 attenuates acute kidney dysfunction and injury markers induced by renal ischemia/reperfusion (I/R) in mice. **A:** The administration of various concentrations of **RNB-61** alleviated I/R induced plasma creatinine levels comparable to the positive control, Fenoldopam (20 mg kg ^-1^) (n=24,12, 10,10,12,17 for Vehicle, Fenoldopam, 0.1, 0.3, 3, 30 mg kg^-1^ **RNB-61**, respectively). Statistical significance was determined using Mann-Whitney tests with Bonferroni correction. No significant difference was observed at 0.1 mg kg^-1^ dose for **RNB-61**. **B:** For the same animals as depicted in panel B, plasma BUN levels were measured. **C-E:** For the same animals, three AKI biomarkers was also quantified (NGAL, Osteopontin, KIM-1).

**Figure 6:**
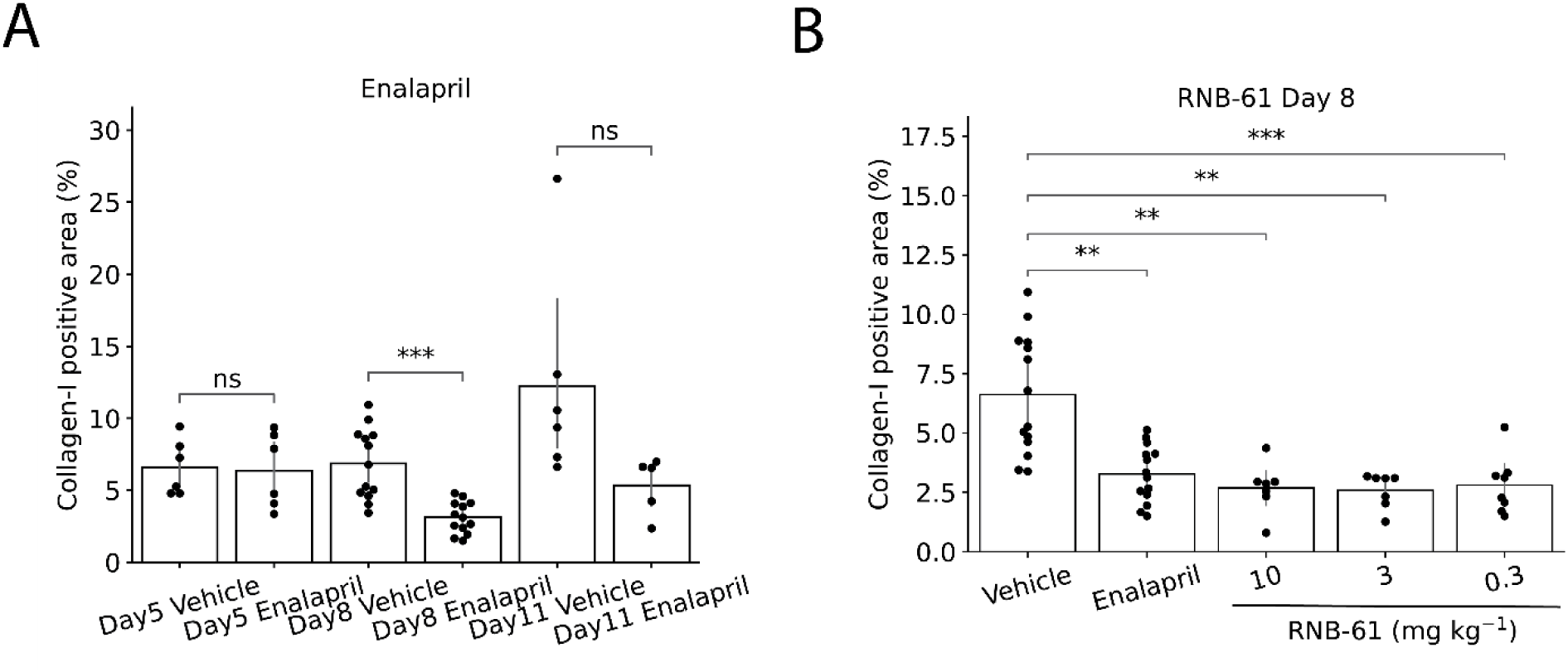
RNB-61 exerts nephroprotective effects in a ureteral obstruction (UOO) rat model of renal fibrosis. **A:** Collagen-III-I positive area was assessed at three consecutive timepoints (d5, d8 and d11) for vehicle controls and rats administered with 32 mg kg^-1^ of enalapril (n=6, 13 and 6 for d5, d8 and d11, respectively). No significant difference was observed at d5 and d11, whereas a significant reduction in the collagen-I positive area was observed at d8. B: The collagen-III-I positive area was assessed after day 8 of the UUO in rats. A significant reduction was observed for Enalapril (32 mg kg^-1^) and for all three doses of **RNB-61** (n=14, 14, 7, 7, 7 for vehicle, enalapril, 0.3, 3 and 3 mg kg^-1^ **RNB-61**, respectively, Mann-Whitney test).

## Discussion

Although the CB2R has been validated as drug target in numerous preclinical models, translation into effective therapeutic agents remains slow ^12,51,52^. The opposite effects of CB1R and CB2R activation in numerous disease models and/or pathological conditions (e.g. liver injury/fibrosis, cardiovascular injury/fibrosis, kidney injury/fibrosis, among others) reflect the distinct roles of CB1R versus CB2R activation in various immune cells, including macrophages, Kupffer cells, osteoclasts, and microglia^19,53^. Thus, the use of optimized receptor (and species)-specific small molecular tools is paramount. Selective CB2R activation typically yields immunosuppressive effects, mitigating sterile inflammation and subsequent tissue damage across numerous pathological conditions without exerting psychoactive effects typically associated with CB1R activation. However, it is worth noting that in certain disease contexts/animal models (e.g. where live pathogens are present), CB2R receptor activation might paradoxically exacerbate or instigate tissue injury^6,54,55^. Several puzzling controversies surrounding CB2R biology and expression/target validation may stem from challenges in detecting the CB2R protein (due to the lack of specific antibodies) and the subpar quality of early tool compounds used in preclinical studies, which lacked selectivity, specificity, and had limited bioavailability^56^.

To resolve the apparent contradictions around the therapeutic roles of CB2Rs from mouse models, in addition to conditional tissue-specific knockout mouse lines, selective CB2R receptor ligands with ideal PK properties are essential to determine the physiologically relevant roles of CB2R in health and disease. In a collaborative research effort, we synthesized and profiled a set of highly potent CB2R ligands from literature, including patent literature, here called **RNB** (Roche, HIH and Bern) compounds. **RNB-61** was characterized for the first time in this study. We demonstrate that beyond its remarkable potency and selectivity, **RNB-61** showcases an ideal physicochemical and pharmacokinetic profile. Moreover, its peripherally restricted action enhances its suitability as a premier pharmacological tool for dissecting the pharmacological impacts of CB2R across various mammalian cellular systems and animal models.

### Selectivity and potency

Out of the five distinct CB2R agonist evaluated in the current report, **RNB-61** showed the highest (∼7,000 fold) selectivity towards *h*CB2 against *h*CB1 and show no significant interaction (defined as >50% inhibition) for 80 additional receptor targets up to 1 µM in a CEREP screen. At 10 µM, which is well above the expected physiological concentration of the compound *in vivo*, the only apparent interaction was with the Na^+^ channel. However, given the peripherally restricted action of **RNB-61**, which was confirmed by measuring P-gp interaction, the effect of **RNB-61** on the Na^+^ ion channel site 2, which is primarily expressed in neurons of the CNS, is unlikely to translate into marked off-target effects in animal models. Another critical aspect of **RNB-61** for the application in relevant animal models is that in addition to *h*CB2R, the binding affinity of **RNB-61** towards *m*CB2R was similar (EC_50_<50 nM). Furthermore, in a functional assay, **RNB-61** also showed comparable potency in four commonly used species for preclinical applications (EC_50_=0.13–1.86 nM). Thus, the excellent selectivity profile of **RNB-61** enabled the specific labeling of CB2Rs in membrane preparations from various immune cell lines as well as *ex vivo* tissue slices, which was also supported by the upregulation of CB2R expression upon the LPS induced inflammatory response in spleen samples of mice.

### Physiochemical and pharmacokinetic properties

As pointed out by Wu et al.^12^ an optimal balance between selectivity, activity, and pharmacokinetic properties of CB2R ligands needs to be achieved. The pyrazole-derived **RNB-61** possesses several favorable physiochemical and pharmacokinetic properties. Unlike typical CBR ligands, **RNB-61** has a logD value of 3.3, indicating moderate lipophilicity which accounts for a good balance between solubility and permeability, essential for optimal oral absorption. In accordance, our measurements in four conditions all indicated that **RNB-61** possesses good aqueous solubility (194, 316, 630 and 1373 µg/mL for LYSA, THESA, FaSSiF and FeSSiF, respectively) that was unexpected due to the high melting point of 188 °C (Table 3). The basic nitrogen atom in the pyrazole ring (N1, basic pKa: 7.37) likely exerts a positive impact on the solubility, which can be exploited for *in vivo* studies (e.g. i.v. or i.p. administration). In terms of permeability, **RNB-61** crossed cell membranes with a relevant effective permeability of 0.9 x 10^-6^ cm s^-1^. In rodent single-dose PK studies, the compound was subject to a low plasma clearance, accounting for ∼5% of hepatic blood flow and in agreement with the intrinsic clearance estimated *in vitro* (Table 3). **RNB-61** displayed an intermediate volume of distribution at steady state (V_ss_= 1.6–2.4 L kg^-1^), suggesting extensive tissue distribution (Figure 4). Taken together, these parameters translated into a terminal plasma half-life of 4–8 h in rats and mice (Table 4).

### Peripherally restricted action

Different peripherally restricted CB2R agonists have been reported, like the AstraZeneca CB2R ligand, AZD194 or the GlaxoSmithKline’s CB2R agonist, GW842166X ^57,58^. Despite the low polar surface of 51 Å^2^, and the presence of only one hydrogen bond donor, **RNB-61** is a strong P-gp substrate in both human (40.6) and mouse (26.8), thus hindering its accumulation in effective concentrations in the CNS. Considering the role of CB2R in brain inflammation, a peripherally restricted CB2R full agonist like **RNB-61** may be useful to address the role of the CB2R in immune cell infiltration versus microglial activation in brain. Given the poor selectivity of several CB2R ligands over CB1R, pharmacological experiments addressing the role of CB2Rs in the brain (a topic that is under scientific debate) could potentially be confounded by CB1R agonism. To elucidate the roles of CB2R in neuroinflammatory and neurodegenerative diseases, the co-administration of **RNB-61** and the P-gp inhibitor tariquidar could be performed, mitigating the ambiguities arising from CB1R activity^41^ (Table 5).

### Nephroprotective effects

The nephroprotective effects of CB2R signaling have been described using acute kidney injury (AKI), which can often progress to chronic kidney disease (CKD), a debilitating condition affecting more than 10% of the global population. This progressive ailment culminates in kidney fibrosis and failure, presenting significant treatment challenges that remain largely unaddressed. Given the wealth of recent studies highlighting the protective role of CB2R signaling and synthetic agonists in diverse preclinical models of acute and chronic kidney diseases^59^, including those induced by the chemotherapy drug cisplatin^20,21,60^, advanced liver injury (hepatorenal syndrome)^22^, chronic diabetes^15,29^, I/R^23,24^, and UUO^25,26^, we investigated the efficacy of **RNB-61** in the I/R-induced model of AKI in mice and the UUO-induced kidney fibrosis model in rats. Consistently with previous studies bilateral kidney I/R injury was associated with significant elevations of serum markers of kidney dysfunction (BUN and creatinine) and parenchymal injury (NGAL, osteopontin, and KIM-1). In the rat model progressive renal fibrosis developed within 8 days following UUO. **RNB-61** exerted dose-dependent tissue protective and/or antifibrotic effects in both mice and rats, which were comparable to the effects of the corresponding reference compounds (Fenoldopam, Tempol, Enalapril).

## Conclusions

In summary, our data show that **RNB-61** is a highly potent and bioavailable CB2R-selective full agonist, which serves as optimal tool compound to investigate the pharmacology of CB2R activation *in vitro* and *in vivo*. Being a substrate for P-gp, **RNB-61** can be used either as peripherally restricted CB2R agonist or CNS penetrating compound if co-administered with a P-gp inhibitor, allowing the differential investigation of the roles of CB2Rs in the periphery and the CNS, without interfering with CB1R activity. In addition, our results support the therapeutic potential of CB2R agonists to treat acute and/or chronic kidney diseases.

## Acknowledgements

The authors wish to thank Camille Perret, Jonathan Mochel, and Franz Schuler for supporting the conduct of the PK and ADME experiments.

## Data availability statement

Raw data supporting the conclusions of this article will be made available by the authors upon request.

## Author contributions (CRediT)

**ACh:** Conceptualization, Investigation, Methodology, Validation, Data Curation, Writing – Original Draft; **DB:** Software, Formal analysis, Visualization, Writing – Original Draft; **CU:** Methodology, Resources, Writing – Original Draft; **ACa:** Methodology, Writing – Review & Editing; **JF:** Methodology, Investigation, Resources, Writing – Review & Editing; **TH:** Methodology, Supervision, Writing – Review & Editing; **RD:** Investigation, Methodology, Writing – Review & Editing; **MM:** Investigation, Methodology, Writing - Review & Editing; **UG:** Investigation, Methodology, Writing – Original Draft, Supervision; **PP:** Conceptualization, Resources, Writing-Original Draft, Supervision; **JG:** Conceptualization, Resources, Supervision, Writing-Original Draft, Review & Editing.

## Supplementary information

**Scheme S1:**
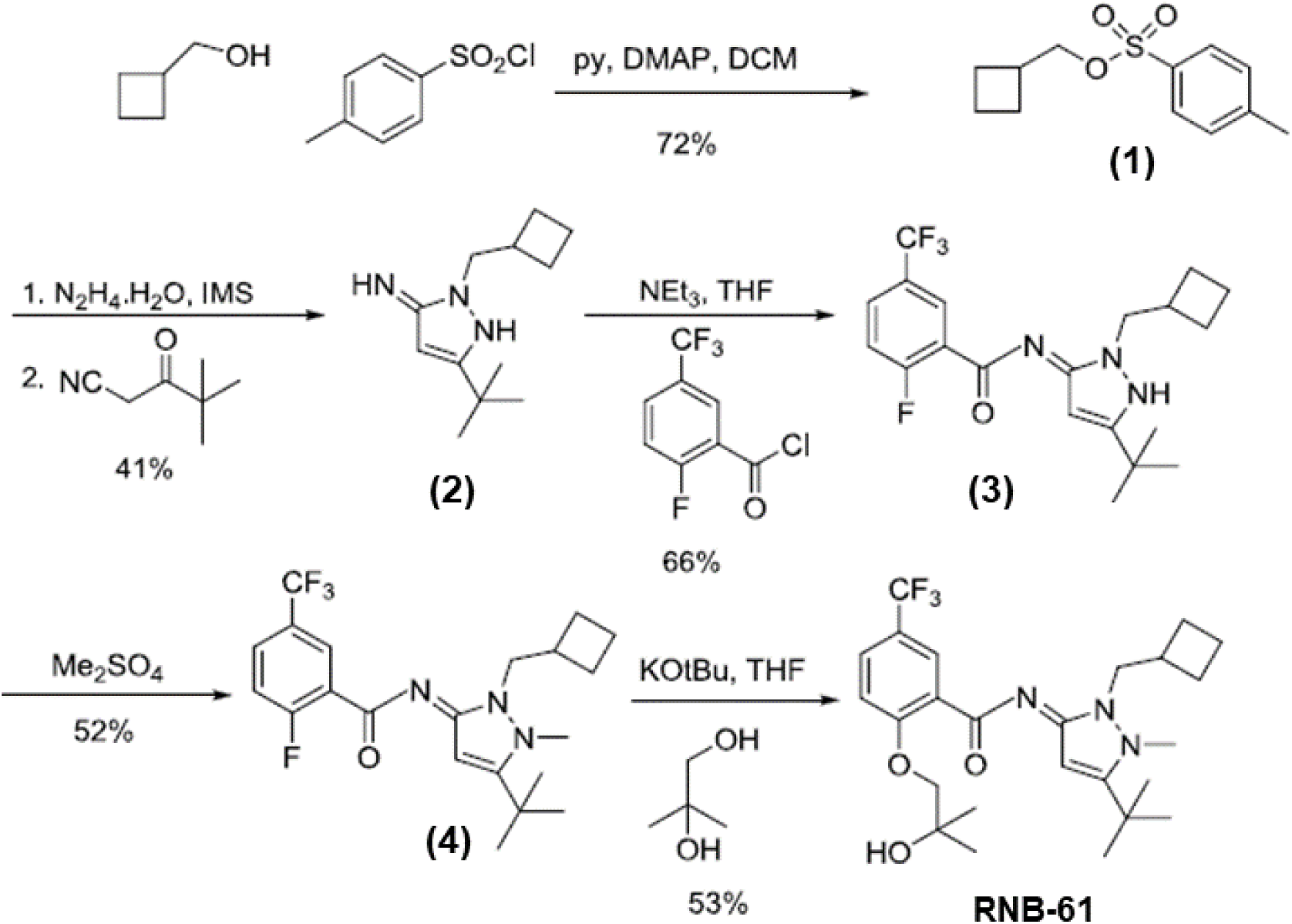
Synthesis of RNB-61.

### Detailed synthesis procedures

Preparation of N-(5-tert-Butyl-2-(cyclobutylmethyl)-1-methyl-1H-pyrazol-3(2H)-ylidene)-2-(2-hydroxy-2-methylpropoxy)-5-(trifluoromethyl)benzamide (**RNB-61**)

*Preparation of 1-(2-Cyclobutylethylsulphonyl)-4-methylbenzene (**1**):* To a solution of cyclobutylmethanol (150 g, 1.74 mol) and pyridine (600 mL) in dichloromethane (1.5 l) was added p-toluenesulphonyl chloride (332 g, 1.74 mol). The solution was allowed to stand overnight and was then treated with 5% hydrochloric acid (180 mL). The organic layer was separated, dried and concentrated. Dry flash chromatography (ethyl acetate → heptane) gave **(1)** as a colourless oil (303 g, 72%).

*Preparation of 5-tert-Butyl-2-(cyclobutylmethyl)-2,3-dihydro-1H-pyrazol-3-amine (**2**):* A mixture of **(1)** (303 g, 1.26 mol), hydrazine hydrate (93 g, 1.9 mol) and IMS (1.5 L) was heated at reflux for 8 h, cooled to room temperature, treated with trimethylacetylacetonitrile (237 g, 1.9 mol) and heated at reflux for a further 5 h. On cooling the solution was concentrated and treated with dichloromethane (3 L). The resultant solution was stirred with saturated aqueous sodium bicarbonate (1.5 L). The organic layer was separated, dried and concentrated to an oil. Dry flash chromatography (ethyl acetate → heptane) gave a solid, which was recrystallised from hexane to give **(2)** as an off-white solid (108 g, 41%).

*Preparation of N-(5-tert-Butyl-2-(cyclobutylmethyl)-1H-pyrazol-3(2H)-ylidene)-2-fluoro-5-(trifluoromethyl)benzamide (**3**):* To a solution of **(2)** (108 g, 0.52 mol) and triethylamine (210g, 2.1 mol) in THF (3 L) was added dropwise over 0.5 h to a solution of 2-fluoro-5-trifluoromethylbenzoyl chloride (118 g, 0.52 mol) in THF (500 mL). On stirring for a further 0.5 h, water (4 L) was added and the product was extracted into dichloromethane (2 × 1500 mL), washed with water (2 × 1 L), dried and concentrated to an oil. Dry flash chromatography (ethyl acetate → heptane) gave **(3)** as a white solid (130 g, 66%).

*Preparation of N-(5-tert-Butyl-2-(cyclobutylmethyl)-1-methylpyrazol-3(2H)-ylidene)-2-fluoro-5-(trifluoromethyl)benzamide (**4**):* **(3)** (130 g, 0.33 mol) and dimethyl sulphate (1.3 L) were stirred at 100°C for 24 h, cooled and poured into 5% aqueous ammonia (7 L). The product was extracted into ethyl acetate (3 × 1 L) and the organic fraction was separated, washed with water (2 L) then brine (500 mL), dried and concentrated. Dry flash chromatography (ethyl acetate → heptane) gave **(4)** as an oil (71 g, 52%) that solidified upon standing.

*Preparation of N-(5-tert-Butyl-2-(cyclobutylmethyl)-1-methyl-1H-pyrazol-3(2H)-ylidene)-2-(2-hydroxy-2-methylpropoxy)-5-(trifluoromethyl)benzamide (**RNB-61**):* To a solution of 2-methylpropane-1,2-diol (34 g, 0.38 mol) in THF (2 L) was added potassium tert-butoxide (39 g, 0.35 mol). On stirring for 0.5 h a solution of **(4)** (71 g, 0.173 mol) in THF (500 mL) was added. The mixture was stirred overnight then poured into water (2 L). The product was extracted into ethyl acetate (2 × 1500 mL), washed with water (2 x 2 L), dried and concentrated to an oil. Recrystallisation from toluene gave **RNB-61** as a white solid (45 g, 53%), mp 191-192°C (eff). The reaction of the amide with dimethyl sulfate in the presence of potassium carbonate led selectively to amide nitrogen methylation yielding the inactive regioisomer of **RNB-61**.

---

**Figure S1:**
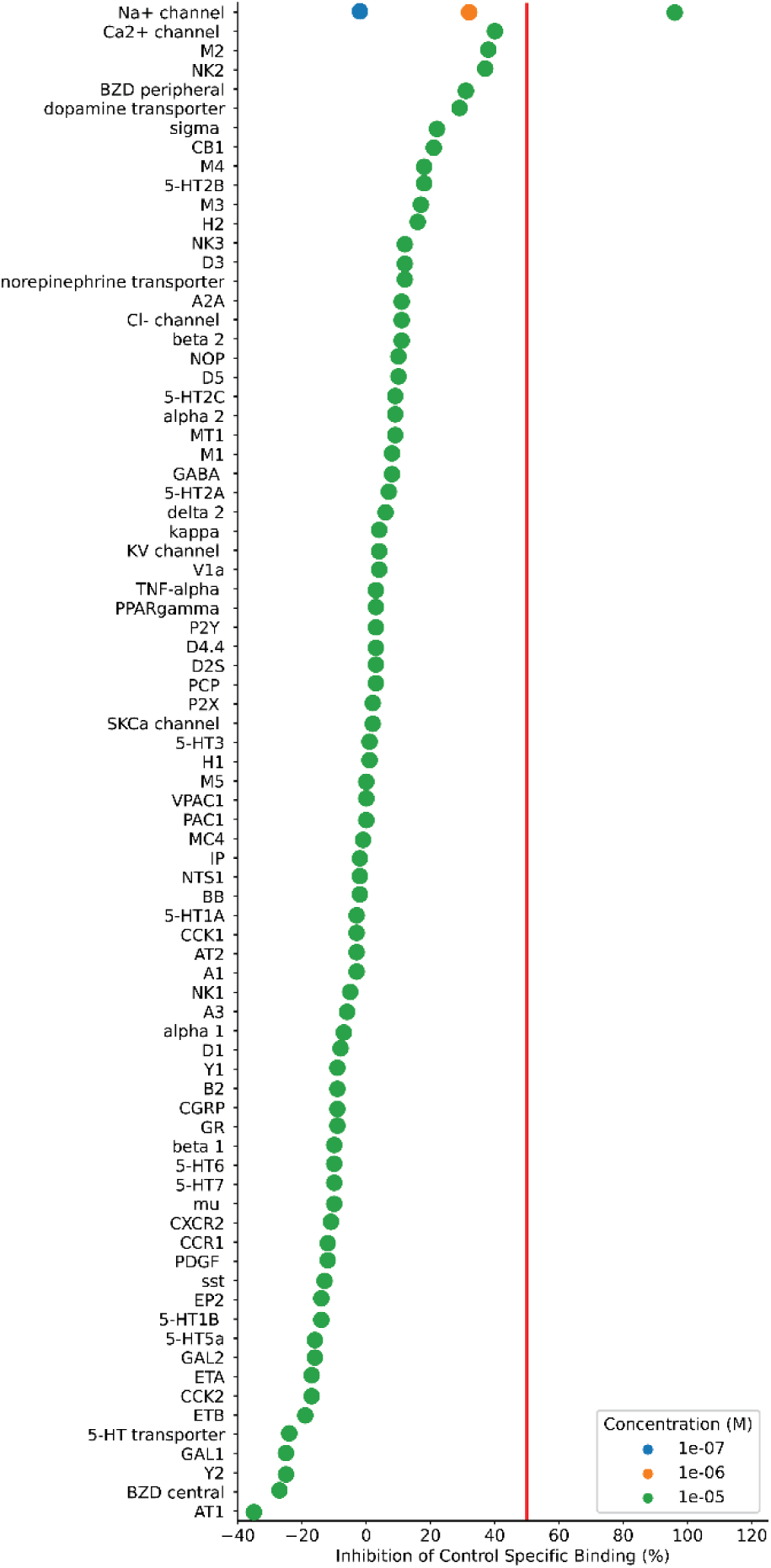
CEREP screening for off-targets. The plot depicts the binding affinity of 0.1-10 µM of **RNB-61**, expressed as the percentage of inhibition of control specific binding on 80 receptors. The threshold for follow-up was set to 50% at 10 µM concentration. Only the Na+ channel showed strong RNB-61 binding (96%) at 10 M, therefore, a follow-up of 1 µM and 0.1 µM was conducted, which showed 32% and 2% inhibition, respectively.

**Table S1:**
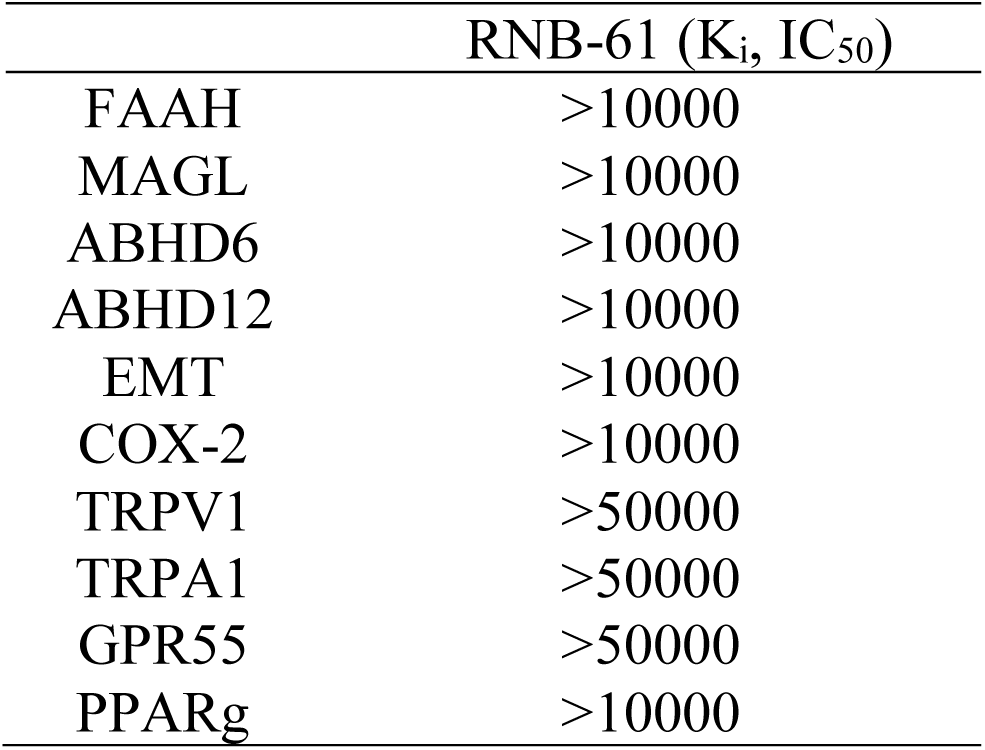
RNB-61 shows no binding interaction and functional inhibition on endocannabinoid related target proteins. Abbreviations: FAAH, Fatty-acid amide hydrolase 1; MAGL, Monoacylglycerol lipase; ABHD, alpha/beta-Hydrolase domain containing; EMT, Endocannabinoid membrane transporter; COX-2, Cyclooxygenase-2; TRPV1, Transient receptor potential vanilloid 1; TRPA1, Transient receptor potential cation channel, subfamily A, member 1; GPR55, G protein-coupled receptor 55, PPARG, Peroxisome Proliferator Activated Receptor Gamma

## References

1. Zou S, Kumar U. Cannabinoid Receptors and the Endocannabinoid System: Signaling and Function in the Central Nervous System. International Journal of Molecular Sciences. 2018;19(3). Available at: https://www.ncbi.nlm.nih.gov/pmc/articles/PMC5877694/.

2. Maccarrone M, Di Marzo V, Gertsch J, et al. Goods and Bads of the Endocannabinoid System as a Therapeutic Target: Lessons Learned after 30 Years. Pharmacological reviews. 2023;75(5):885–958. Available at: https://pubmed.ncbi.nlm.nih.gov/37164640/.

3. Bie B, Wu J, Foss JF, Naguib M. An overview of the cannabinoid type 2 receptor system and its therapeutic potential. Current opinion in anaesthesiology. 2018;31(4):407–414. Available at: https://www.ncbi.nlm.nih.gov/pmc/articles/PMC6035094/.

4. Dhopeshwarkar A, Mackie K. CB2 Cannabinoid receptors as a therapeutic target-what does the future hold? Molecular Pharmacology. 2014;86(4):430–437. Available at: https://www.ncbi.nlm.nih.gov/pmc/articles/PMC4164977/.

5. Steffens S, Pacher P. Targeting cannabinoid receptor CB(2) in cardiovascular disorders: promises and controversies. British Journal of Pharmacology. 2012;167(2):313–323. Available at: https://pubmed.ncbi.nlm.nih.gov/22612332/.

6. Pacher P, Mechoulam R. Is lipid signaling through cannabinoid 2 receptors part of a protective system? Progress in lipid research. 2011;50(2):193–211. Available at: https://www.ncbi.nlm.nih.gov/pmc/articles/PMC3062638/.

7. Devane WA, Hanus L, Breuer A, et al. Isolation and structure of a brain constituent that binds to the cannabinoid receptor. *Science (New York*, N.Y.). 1992;258(5090):1946-1949. Available at: https://pubmed.ncbi.nlm.nih.gov/1470919/.

8. Mechoulam R, Ben-Shabat S, Hanus L, et al. Identification of an endogenous 2-monoglyceride, present in canine gut, that binds to cannabinoid receptors. Biochemical pharmacology. 1995;50(1):83–90. Available at: https://pubmed.ncbi.nlm.nih.gov/7605349/.

9. Banister SD, Connor M. The Chemistry and Pharmacology of Synthetic Cannabinoid Receptor Agonist New Psychoactive Substances: Evolution. Handbook of experimental pharmacology. 2018;252:191–226. Available at: https://pubmed.ncbi.nlm.nih.gov/30105473/.

10. Di Marzo V. The endocannabinoid system: its general strategy of action, tools for its pharmacological manipulation and potential therapeutic exploitation. Pharmacological research. 2009;60(2):77–84. Available at: https://pubmed.ncbi.nlm.nih.gov/19559360/.

11. Gasperi V, Guzzo T, Topai A, et al. Recent Advances on Type-2 Cannabinoid (CB2) Receptor Agonists and their Therapeutic Potential. Current medicinal chemistry. 2023;30(12):1420–1457. Available at: https://pubmed.ncbi.nlm.nih.gov/36028971/.

12. Wu Y-R, Tang J-Q, Zhang W-N, Zhuang C-L, Shi Y. Rational drug design of CB2 receptor ligands: from 2012 to 2021. RSC Adv. 2022;12(54):35242–35259. Available at: https://pubs.rsc.org/en/content/articlelanding/2022/ra/d2ra05661e.

13. Starowicz K, Finn DP. Cannabinoids and Pain: Sites and Mechanisms of Action. Advances in pharmacology (San Diego, Calif.). 2017;80:437–475. Available at: https://pubmed.ncbi.nlm.nih.gov/28826543/.

14. Yoo EH, Lee JH. Cannabinoids and Their Receptors in Skin Diseases. International Journal of Molecular Sciences. 2023;24(22):16523. Available at: https://www.mdpi.com/1422-0067/24/22/16523.

15. Barutta F, Piscitelli F, Pinach S, et al. Protective role of cannabinoid receptor type 2 in a mouse model of diabetic nephropathy. Diabetes. 2011;60(9):2386–2396. Available at: https://pubmed.ncbi.nlm.nih.gov/21810593/.

16. Lotersztajn S, Teixeira-Clerc F, Julien B, et al. CB2 receptors as new therapeutic targets for liver diseases. British Journal of Pharmacology. 2008;153(2):286–289. Available at: https://www.ncbi.nlm.nih.gov/pmc/articles/PMC2219531/.

17. Bátkai S, Osei-Hyiaman D, Pan H, et al. Cannabinoid-2 receptor mediates protection against hepatic ischemia/reperfusion injury. FASEB journal : official publication of the Federation of American Societies for Experimental Biology. 2007;21(8):1788–1800. Available at: https://pubmed.ncbi.nlm.nih.gov/17327359/.

18. Pacher P, Haskó G. Endocannabinoids and cannabinoid receptors in ischaemia-reperfusion injury and preconditioning. British Journal of Pharmacology. 2008;153(2):252–262. Available at: https://pubmed.ncbi.nlm.nih.gov/18026124/.

19. Gertsch J. Editorial: Lung macrophages high on cannabinoids: jamming PAMs and taming TAMs? Journal of leukocyte biology. 2016;99(4):518–520. Available at: https://pubmed.ncbi.nlm.nih.gov/27034462/.

20. Mukhopadhyay P, Rajesh M, Pan H, et al. Cannabinoid-2 receptor limits inflammation, oxidative/nitrosative stress, and cell death in nephropathy. Free radical biology & medicine. 2010;48(3):457–467. Available at: https://pubmed.ncbi.nlm.nih.gov/19969072/.

21. Mukhopadhyay P, Baggelaar M, Erdelyi K, et al. The novel, orally available and peripherally restricted selective cannabinoid CB2 receptor agonist LEI-101 prevents cisplatin-induced nephrotoxicity. British Journal of Pharmacology. 2016;173(3):446–458. Available at: https://www.ncbi.nlm.nih.gov/pmc/articles/PMC4728411/.

22. Trojnar E, Erdelyi K, Matyas C, et al. Cannabinoid-2 receptor activation ameliorates hepatorenal syndrome. Free radical biology & medicine. 2020;152:540–550. Available at: https://www.sciencedirect.com/science/article/pii/S0891584919317459.

23. Çakır M, Tekin S, Doğanyiğit Z, Çakan P, Kaymak E. The protective effect of cannabinoid type 2 receptor activation on renal ischemia-reperfusion injury. Molecular and cellular biochemistry. 2019;462(1-2):123–132. Available at: https://pubmed.ncbi.nlm.nih.gov/31446615/.

24. Pressly JD, Mustafa SM, Adibi AH, et al. Selective Cannabinoid 2 Receptor Stimulation Reduces Tubular Epithelial Cell Damage after Renal Ischemia-Reperfusion Injury. The Journal of pharmacology and experimental therapeutics. 2018;364(2):287–299. Available at: https://pubmed.ncbi.nlm.nih.gov/29187590/.

25. Nettekoven M, Adam J-M, Bendels S, et al. Novel Triazolopyrimidine-Derived Cannabinoid Receptor 2 Agonists as Potential Treatment for Inflammatory Kidney Diseases. ChemMedChem. 2016;11(2):179–189. Available at: https://pubmed.ncbi.nlm.nih.gov/26228928/.

26. Swanson ML, Regner KR, Moore BM, Park F. Cannabinoid Type 2 Receptor Activation Reduces the Progression of Kidney Fibrosis Using a Mouse Model of Unilateral Ureteral Obstruction. Cannabis and cannabinoid research. 2022;7(6):790–803. Available at: https://pubmed.ncbi.nlm.nih.gov/35196117/.

27. Hickey ER, Zindell R, Cirillo PF, et al. Selective CB2 receptor agonists. Part 1: the identification of novel ligands through computer-aided drug design (CADD) approaches. Bioorganic & medicinal chemistry letters. 2015;25(3):575–580. Available at: https://pubmed.ncbi.nlm.nih.gov/25556098/.

28. Long C, Xie N, Shu Y, et al. Knockout of the Cannabinoid Receptor 2 Gene Promotes Inflammation and Hepatic Stellate Cell Activation by Promoting A20/Nuclear Factor-κB (NF-κB) Expression in Mice with Carbon Tetrachloride-Induced Liver Fibrosis. Medical Science Monitor : International Medical Journal of Experimental and Clinical Research. 2021;27:e931236. Available at: https://www.ncbi.nlm.nih.gov/pmc/articles/PMC8409143/.

29. Barutta F, Grimaldi S, Franco I, et al. Deficiency of cannabinoid receptor of type 2 worsens renal functional and structural abnormalities in streptozotocin-induced diabetic mice. Kidney international. 2014;86(5):979–990. Available at: https://pubmed.ncbi.nlm.nih.gov/24827776/.

30. Soethoudt M, Grether U, Fingerle J, et al. Cannabinoid CB2 receptor ligand profiling reveals biased signalling and off-target activity. Nat Commun. 2017;8(1):13958. Available at: https://www.nature.com/articles/ncomms13958#Sec12.

31. Yao BB, Hsieh GC, Frost JM, et al. In vitro and in vivo characterization of A-796260: a selective cannabinoid CB2 receptor agonist exhibiting analgesic activity in rodent pain models. British Journal of Pharmacology. 2008;153(2):390–401. Available at: https://www.ncbi.nlm.nih.gov/pmc/articles/PMC2219533/.

32. William A. Carroll, Michael J. Dart, Jennifer M. Frost, Steven P. Latshaw, Teodozyj Kolasa, Tongmei Li, Sridhar Peddi, Bo Liu, Arturo Perez-Medrano, Meena Patel, Xueqing Wang, Derek W. Nelson, inventor. Novel compounds as cannabinoid receptor ligands. US20100069348A1.

33. Cheng Y, Albrecht BK, Brown J, et al. Discovery and optimization of a novel series of N-arylamide oxadiazoles as potent, highly selective and orally bioavailable cannabinoid receptor 2 (CB2) agonists. Journal of medicinal chemistry. 2008;51(16):5019–5034. Available at: https://pubmed.ncbi.nlm.nih.gov/18680277/.

34. Pier Francesco Cirillo, Eugene Richard Hickey, Doris Riether, Monika Ermann, Innocent Mushi, inventor; Boehringer Ingelheim International GMBH. Amine and ether compounds which modulate the cb2 receptor. WO2009105509A1.

35. Alessandra Bartolozzi, Eugene Richard Hickey, Doris Riether, Lifen Wu, Renee M. Zindell, Stephen Peter East, Monika Ermann, inventor; Boehringer Ingelheim International GMBH. Compounds Which Selectively Modulate The CB2 Receptor. US20100076029A1.

36. Chicca A, Nicolussi S, Bartholomäus R, et al. Chemical probes to potently and selectively inhibit endocannabinoid cellular reuptake. Proc. Natl. Acad. Sci. U.S.A. 2017;114(25):E5006–E5015. Available at: https://www.pnas.org/doi/full/10.1073/pnas.1704065114.

37. Porter RF, Szczesniak A-M, Toguri JT, et al. Selective Cannabinoid 2 Receptor Agonists as Potential Therapeutic Drugs for the Treatment of Endotoxin-Induced Uveitis. Molecules (Basel, Switzerland). 2019;24(18). Available at: https://pubmed.ncbi.nlm.nih.gov/31540271/.

38. Wagner B, Fischer H, Kansy M, Seelig A, Assmus F. Carrier Mediated Distribution System (CAMDIS): a new approach for the measurement of octanol/water distribution coefficients. European journal of pharmaceutical sciences : official journal of the European Federation for Pharmaceutical Sciences. 2015;68:68–77. Available at: https://pubmed.ncbi.nlm.nih.gov/25513709/.

39. Mock ED, Mustafa M, Gunduz-Cinar O, et al. Discovery of a NAPE-PLD inhibitor that modulates emotional behavior in mice. Nat Chem Biol. 2020;16(6):667–675. Available at: https://www.nature.com/articles/s41589-020-0528-7.

40. He Y, Schild M, Grether U, et al. Development of High Brain-Penetrant and Reversible Monoacylglycerol Lipase PET Tracers for Neuroimaging. Journal of medicinal chemistry. 2022;65(3):2191–2207. Available at: https://www.dora.lib4ri.ch/psi/islandora/object/psi%3A40924.

41. Fowler S, Zhang H. In vitro evaluation of reversible and irreversible cytochrome P450 inhibition: current status on methodologies and their utility for predicting drug-drug interactions. The AAPS journal. 2008;10(2):410–424. Available at: https://pubmed.ncbi.nlm.nih.gov/18686042/.

42. Brink A, Fontaine F, Marschmann M, et al. Post-acquisition analysis of untargeted accurate mass quadrupole time-of-flight MS(E) data for multiple collision-induced neutral losses and fragment ions of glutathione conjugates. Rapid communications in mass spectrometry : RCM. 2014;28(24):2695–2703. Available at: https://pubmed.ncbi.nlm.nih.gov/25380491/.

43. Zhang W, Guo L, Liu H, et al. Discovery of Linvencorvir (RG7907), a Hepatitis B Virus Core Protein Allosteric Modulator, for the Treatment of Chronic HBV Infection. Journal of medicinal chemistry. 2023;66(6):4253–4270.

44. Haider A, Gobbi L, Kretz J, et al. Identification and Preclinical Development of a 2,5,6-Trisubstituted Fluorinated Pyridine Derivative as a Radioligand for the Positron Emission Tomography Imaging of Cannabinoid Type 2 Receptors. Journal of medicinal chemistry. 2020;63(18):10287–10306.

45. Poirier A, Cascais A-C, Bader U, et al. Calibration of in vitro multidrug resistance protein 1 substrate and inhibition assays as a basis to support the prediction of clinically relevant interactions in vivo. Drug metabolism and disposition: the biological fate of chemicals. 2014;42(9):1411–1422. Available at: https://pubmed.ncbi.nlm.nih.gov/24939652/.

46. Fox E, Bates SE. Tariquidar (XR9576): a P-glycoprotein drug efflux pump inhibitor. Expert review of anticancer therapy. 2007;7(4):447–459. Available at: https://pubmed.ncbi.nlm.nih.gov/17428165/.

47. Skrypnyk NI, Harris RC, Caestecker MP de. Ischemia-reperfusion model of acute kidney injury and post injury fibrosis in mice. Journal of visualized experiments: JoVE. 2013(78). Available at: https://pubmed.ncbi.nlm.nih.gov/23963468/.

48. Feizi A, Jafari M-R, Hamedivafa F, Tabrizian P, Djahanguiri B. The preventive effect of cannabinoids on reperfusion-induced ischemia of mouse kidney. Experimental and toxicologic pathology : official journal of the Gesellschaft fur Toxikologische Pathologie. 2008;60(4-5):405–410. Available at: https://pubmed.ncbi.nlm.nih.gov/18571910/.

49. Chevalier RL, Forbes MS, Thornhill BA. Ureteral obstruction as a model of renal interstitial fibrosis and obstructive nephropathy. Kidney international. 2009;75(11):1145–1152. Available at: https://pubmed.ncbi.nlm.nih.gov/19340094/.

50. Frost JM, Dart MJ, Tietje KR, et al. Indol-3-ylcycloalkyl ketones: effects of N1 substituted indole side chain variations on CB(2) cannabinoid receptor activity. Journal of medicinal chemistry. 2010;53(1):295–315. Available at: https://pubmed.ncbi.nlm.nih.gov/19921781/.

51. Atwood BK, Straiker A, Mackie K. CB₂: therapeutic target-in-waiting. Progress in neuro-psychopharmacology & biological psychiatry. 2012;38(1):16–20. Available at: https://www.ncbi.nlm.nih.gov/pmc/articles/PMC3345167/.

52. Cabañero D, Martín-García E, Maldonado R. The CB2 cannabinoid receptor as a therapeutic target in the central nervous system. Expert opinion on therapeutic targets. 2021;25(8):659–676. Available at: https://pubmed.ncbi.nlm.nih.gov/34424117/.

53. Horváth B, Magid L, Mukhopadhyay P, et al. A new cannabinoid CB2 receptor agonist HU-910 attenuates oxidative stress, inflammation and cell death associated with hepatic ischaemia/reperfusion injury. British Journal of Pharmacology. 2012;165(8):2462–2478. Available at: https://www.ncbi.nlm.nih.gov/pmc/articles/PMC3423243/.

54. Alferink J, Specht S, Arends H, et al. Cannabinoid Receptor 2 Modulates Susceptibility to Experimental Cerebral Malaria through a CCL17-dependent Mechanism. The Journal of biological chemistry. 2016;291(37):19517–19531. Available at: https://pubmed.ncbi.nlm.nih.gov/27474745/.

55. Maccarrone M, Bab I, Bíró T, et al. Endocannabinoid signaling at the periphery: 50 years after THC. Trends in pharmacological sciences. 2015;36(5):277–296. Available at: https://pubmed.ncbi.nlm.nih.gov/25796370/.

56. Uwe Grether PP. Challenges bringing CB₂R medicine to bedside. Open Access Government. 2023. 01. 03.

57. Naguib M, Diaz P, Xu JJ, et al. MDA7: a novel selective agonist for CB2 receptors that prevents allodynia in rat neuropathic pain models. British Journal of Pharmacology. 2008;155(7):1104–1116. Available at: https://www.ncbi.nlm.nih.gov/pmc/articles/PMC2597252/.

58. Battista N, Di Tommaso M, Bari M, Maccarrone M. The endocannabinoid system: an overview. Frontiers in Behavioral Neuroscience. 2012;6:9. Available at: https://www.ncbi.nlm.nih.gov/pmc/articles/PMC3303140/.

59. Zhao Z, Yan Q, Xie J, et al. The intervention of cannabinoid receptor in chronic and acute kidney disease animal models: a systematic review and meta-analysis. Diabetology & metabolic syndrome. 2024;16(1):45. Available at: https://pubmed.ncbi.nlm.nih.gov/38360685/.

60. Li X, Chang H, Bouma J, et al. Structural basis of selective cannabinoid CB2 receptor activation. Nat Commun. 2023;14(1):1447. Available at: https://pubmed.ncbi.nlm.nih.gov/36922494/.

